# ToDD: Topological Compound Fingerprinting in Computer-Aided Drug Discovery

**DOI:** 10.1101/2022.11.08.515685

**Authors:** Andac Demir, Baris Coskunuzer, Ignacio Segovia-Dominguez, Yuzhou Chen, Yulia Gel, Bulent Kiziltan

**Author notes:** Equal contribution (Andac Demir, Baris Coskunuzer). 36th Conference on Neural Information Processing Systems (*N*eurIPS 2022).

## Abstract

In computer-aided drug discovery (CADD), virtual screening (VS) is used for identifying the drug candidates that are most likely to bind to a molecular target in a large library of compounds. Most VS methods to date have focused on using canonical compound representations (e.g., SMILES strings, Morgan fingerprints) or generating alternative fingerprints of the compounds by training progressively more complex variational autoencoders (VAEs) and graph neural networks (GNNs). Although VAEs and GNNs led to significant improvements in VS performance, these methods suffer from reduced performance when scaling to large virtual compound datasets. The performance of these methods has shown only incremental improvements in the past few years. To address this problem, we developed a novel method using multiparameter persistence (MP) homology that produces topological fingerprints of the compounds as multidimensional vectors. Our primary contribution is framing the VS process as a new topology-based graph ranking problem by partitioning a compound into chemical substructures informed by the periodic properties of its atoms and extracting their persistent homology features at multiple resolution levels. We show that the margin loss fine-tuning of pretrained Triplet networks attains highly competitive results in differentiating between compounds in the embedding space and ranking their likelihood of becoming effective drug candidates. We further establish theoretical guarantees for the stability properties of our proposed MP signatures, and demonstrate that our models, enhanced by the MP signatures, outperform state-of-the-art methods on benchmark datasets by a wide and highly statistically significant margin (e.g., 93% gain for Cleves-Jain and 54% gain for DUD-E Diverse dataset).

## 1 Introduction

Drug discovery is the early phase of the pharmaceutical R&D pipeline where machine learning (ML) is making a paradigm-shifting impact [31, 91]. Traditionally, early phases of biomedical research involve the identification of targets for a disease of interest, followed by high-throughput screening (HTS) experiments to determine hits within the synthesized compound library, i.e., compounds with high potential. Then, these compounds are optimized to increase potency and other desired target properties. In the final phases of the R&D pipeline, drug candidates have to pass a series of rigorous controlled tests in clinical trials to be considered for regulatory approval. On average, this process takes 10-15 years end-to-end and costs in excess of ~ 2 billion US dollars [10]. HTS is highly time and cost-intensive. Therefore, it is critical to find good potential compounds effectively for the HTS step for novel compound discovery, but also to speed up the pipeline and make it more cost-effective. To address this need, ML augmented virtual screening (VS) has emerged as a powerful computational approach to screen ultra large libraries of compounds to find the ones with desired properties and prioritize them for experimentation [65, 40].

In this paper, we develop novel approaches for virtual screening by successfully integrating topological data analysis (TDA) methods with ML and deep learning (DL) tools. We first produce topological fingerprints of compounds as 2*D* or 3*D* vectors by using TDA tools, i.e., multidimensional persistent homology. Then, we show that Triplet networks, (where state-of-the-art pretrained transformer-based models and modernized convolutional neural network architectures serve as the backbone and distinct topological features allow to represent support and query compounds), successfully identify the compounds with the desired properties. We also demonstrate that the applicability of topological feature maps can be successfully generalized to traditional ML algorithms such as random forests.

The distinct advantage of TDA tools, in particular persistent homology (PH), is that it enables effective integration of the domain information such as atomic mass, partial charge, bond type (single, double, triple, aromatic ring), ionization energy or electron affinity, which carry vital information regarding the chemical properties of a compound at multiple resolution levels during the graph filtration step. While common PH theory allows only one such domain function to be used in this process, with our novel multipersistence approach, we show it is possible to use more than one domain function. Topological fingerprints can effectively carry much finer chemical information of the compound structure informed by the multiple domain functions embedded in the process. Specifically, multiparameter persistence homology decomposes a 2*D* graph structure of a compound into a series of subgraphs using domain functions and generates hierarchical topological representations in multiple resolution levels. At each resolution stage, our algorithm sequentially generates finer topological fingerprints of the chemical substructures. We feed these topological fingerprints to suitable ML/DL methods, and our ToDD models achieve state-of-the-art in all benchmark datasets across all targets (See Table 1 and 2).

**Table 1:** Comparison of EF 2%, 5%, 10% and ROC-AUC values between ToDD and other virtual screening methods on the Cleves-Jain dataset.

| Model | EF 2% (max. 50) | EF 5% (max. 20) | EF 10% (max. 10) | ROC-AUC |
| --- | --- | --- | --- | --- |
| USR [8] | 10.0 | 6.2 | 4.1 | 0.76 |
| GZD [92] | 13.4 | 8.0 | 5.3 | 0.81 |
| PS [42] | 10.7 | 6.6 | 4.9 | 0.78 |
| ROCS [38] | 20.1 | 10.7 | 6.2 | <u>0.83</u> |
| USR + GZD [84] | 13.7 | 7.7 | 4.7 | 0.81 |
| USR + PS [84] | 13.1 | 7.9 | 5.0 | 0.80 |
| USR + ROCS [84] | 17.1 | 9.1 | 5.4 | <u>0.83</u> |
| GZD + PS [84] | 16.0 | 9.1 | 5.9 | 0.82 |
| PH_VS [50] | 18.6 | NA | NA | NA |
| GZD + ROCS [84] | 20.3 | <u>10.8</u> | 5.3 | <u>0.83</u> |
| PS + ROCS [84] | <u>20.5</u> | 10.7 | 6.4 | <u>0.83</u> |
| **ToDD-RF** | 35.2±2.3 | 15.6±1.0 | 8.1±0.4 | **0.94±0.02** |
| **ToDD-ViT** | **39.6±1.4** | **18.6±0.4** | **9.9±0.1** | 0.90±0.01 |
| Relative gains | 92.9% | 83.7% | 54.1% | 13.3% |

**Table 2:** Comparison of EF 1% (max. 100) between ToDD and other virtual screening methods on 8 targets of the DUD-E Diverse subset.

| Model | AMPC | CXCR4 | KIF11 | CP3A4 | GCR | AKT1 | HIVRT | HIVPR | Avg. |
| --- | --- | --- | --- | --- | --- | --- | --- | --- | --- |
| Findsite [101] | 0.0 | 0.0 | 0.9 | 21.7 | 34.2 | 39.0 | 1.2 | 34.7 | 16.5 |
| FragSite [102] | 4.2 | 42.5 | 0.0 | 32.9 | 29.1 | 47.1 | 2.4 | 48.7 | 25.9 |
| Gnina [87] | 2.1 | 15.0 | 38.0 | 1.2 | 39.0 | 4.1 | 11.0 | 28.0 | 17.3 |
| GOLD-EATL [96] | 25.8 | 20.0 | 33.5 | 17.9 | 34.6 | 29.2 | 28.7 | 23.4 | 26.6 |
| Glide-EATL [96] | 35.5 | 20.8 | 30.5 | 15.1 | 24.0 | 31.6 | 29.0 | 22.0 | 26.1 |
| CompM [96] | 32.3 | 25.0 | 35.5 | 33.6 | 37.1 | 44.2 | 30.2 | 25.0 | 32.9 |
| CompScore [75] | 39.6 | 51.6 | 51.3 | 14.0 | 27.1 | 37.6 | 21.8 | 18.2 | 32.7 |
| CNN [77] | 2.1 | 5.0 | 11.2 | 28.7 | 12.8 | 84.6 | 12.2 | 9.9 | 20.8 |
| DenseFS [44] | 14.6 | 5.0 | 4.3 | <u>44.3</u> | 20.9 | 89.4 | 12.8 | 8.4 | 25.0 |
| SIEVE-Score [98] | 30.7 | 61.1 | 53.4 | 6.7 | 33.3 | 42.1 | 39.8 | 38.3 | 38.2 |
| DeepScore [94] | 28.1 | 56.8 | <u>54.3</u> | 37.1 | <u>40.9</u> | 59.0 | <u>43.8</u> | 62.8 | <u>47.9</u> |
| RF-Score-VSv3 [98] | 32.3 | 60.9 | 4.5 | 25.9 | 32.5 | 41.9 | 39.8 | <u>65.7</u> | 37.9 |
| **ToDD-RF** | 42.9±4.5 | **92.3±3.2** | **75.0±5.0** | **67.6±3.4** | **78.9±4.0** | **90.7±1.3** | **64.1±2.3** | **92.1±1.5** | **73.7** |
| **ToDD-ConvNeXt** | **46.2±3.6** | 84.6±2.8 | 72.5±3.6 | 28.8±2.8 | 46.0±2.0 | 81.2±2.5 | 37.5±3.6 | 74.6±1.0 | 58.9 |
| Relative gains | 16.7% | 51.1% | 38.1% | 52.6% | 92.9% | 1.5% | 46.3% | 40.2% | 53.9% |

**Table 3:** Summary statistics of the Cleves-Jain dataset.

| Target | # Training Samples | # Test Samples |
| --- | --- | --- |
| a | 3 | 6 |
| b | 3 | 22 |
| c | 2 | 13 |
| d | 3 | 6 |
| e | 2 | 5 |
| f | 2 | 4 |
| g | 2 | 5 |
| h | 2 | 5 |
| i | 2 | 5 |
| j | 3 | 14 |
| k | 3 | 14 |
| l | 3 | 10 |
| m | 3 | 9 |
| n | 2 | 10 |
| o | 3 | 30 |
| p | 3 | 23 |
| q | 3 | 11 |
| r | 2 | 14 |
| s | 3 | 15 |
| t | 2 | 5 |
| u | 3 | 9 |
| v | 3 | 7 |
| Decoy | 0 | 850 |

**Table 4:**
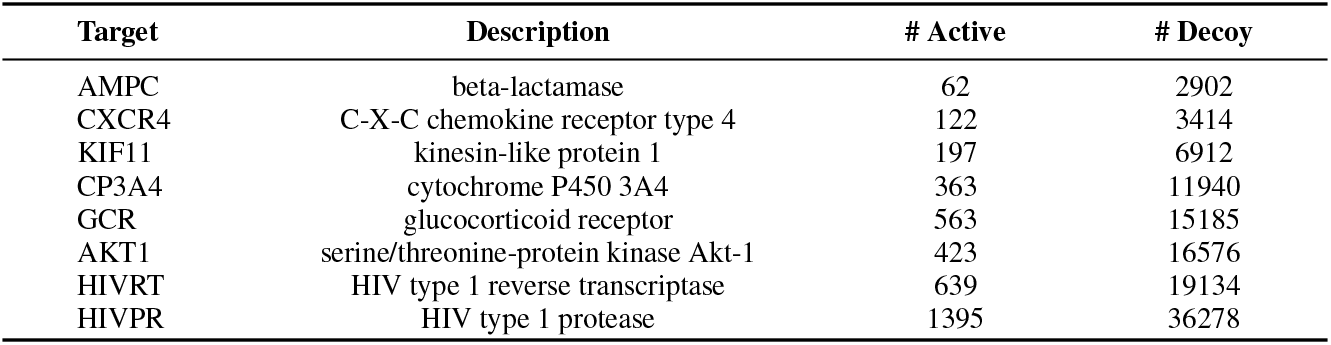
Summary statistics of the DUD-E Diverse dataset.

**The key contributions of this paper are**:

- We develop a transformative approach to generate molecular fingerprints. Using multipersistence, we produce highly expressive and unique topological fingerprints for compounds independent of scale and complexity. This offers a new way to describe and search chemical space relevant to both drug discovery and development.
- We bring a new perspective to multiparameter persistence in TDA and produce a computationally efficient multidimensional fingerprint of chemical data that can successfully incorporate more than one domain function to the PH process. These MP fingerprints harness the computational strength of linear representations and are suitable to be integrated into a broad range of ML, DL, and statistical methods; and open a path for computationally efficient extraction of latent topological information.
- We prove that our multidimensional persistence fingerprints have the same important stability guarantees as the ones exhibited by the most currently existing summaries for single persistence.
- We perform extensive numerical experiments in VS, showing that our ToDD models outperform all state-of-the-art methods by a wide margin (See Figure 1).

**Figure 1:**
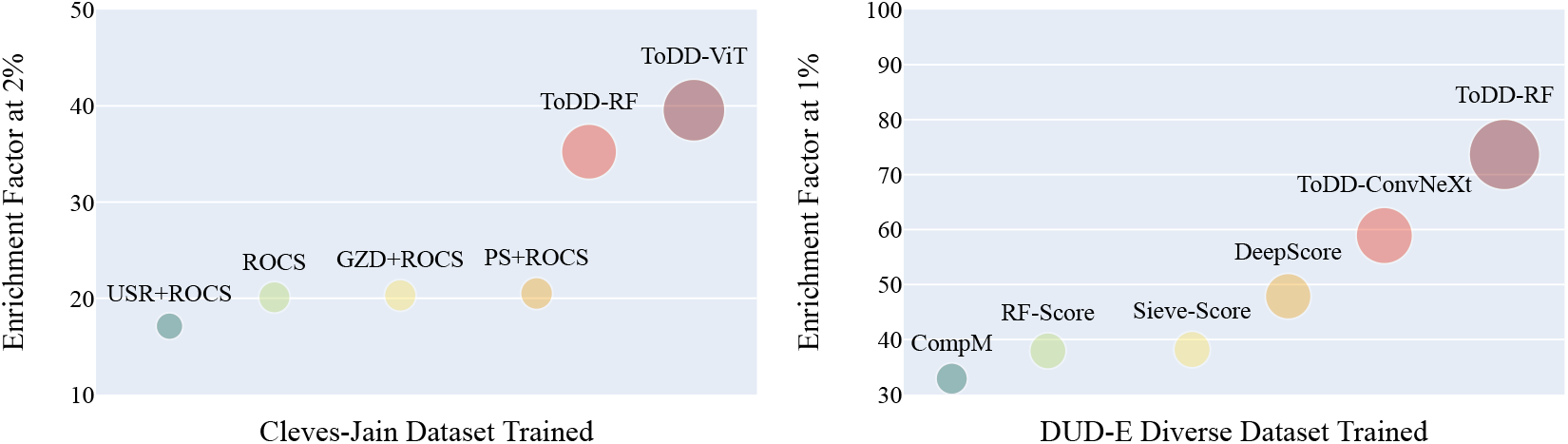
Comparison of virtual screening performance. Each bubble’s diameter is proportional to its EF score. ToDD offers significant gain regardless of the choice of classification model such as random forests (RF), vision transformer (ViT) or a modernized ResNet architecture ConvNeXt. The standard performance metric *EF*_*α*%_ is defined as 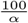, and therefore the maximum attainable value is 50 for *EF*_2%_, and 100 for *EF*_1%_.

## 2 Related Work

### 2.1 Virtual Screening

A key step in the early stages of the drug discovery process is to find active compounds that will be further optimized into potential drug candidates. One prevalent computational method that is widely used for compound prioritization with desired properties is *virtual screening* (VS). There are two major categories, i.e., structure-based virtual screening (SBVS) and ligand-based virtual screening (LBVS) [20]. SBVS uses the 3*D* structural information of both ligand (compound) and target protein as a complex [12, 52, 86]. SBVS methods generally require a good understanding of 3*D*-structure of the target protein to explore the different poses of a compound in a binding pocket of the target. This makes the process computationally expensive. On the other hand, LBVS methods compare structural similarities of a library of compounds with a known active ligand [79, 69] with an underlying assumption that similar compounds are prone to exhibit similar biological activity. Unlike SBVS, LBVS only uses ligand information. The main idea is to produce effective fingerprints of the compounds and use ML tools to find similarities. Therefore, computationally less expensive LBVS methods can be more efficient with larger chemical datasets especially when the structure of the target receptor is not known [56].

In the last 3 decades, various LBVS methods have been developed with different approaches and these can be categorized into 3 classes depending on the fingerprint they produce: SMILES [81] and SMARTS [29] are examples of 1*D*-methods which produce 1*D*-fingerprints, compressing compound information to a vector. RASCAL [78], MOLPRINT2*D* [9], ECFP [80], CDK-graph [97],CDK-hybridization [85],SWISS [103], Klekota-Roth [53], MACSS [29], E-state [36] and SIMCOMP [37] are among 2*D* methods which uses 2*D*-structure fingerprint and graph matching. Finally, examples of 3*D*-methods are ROCS [38], USR [8], PatchSurfer [42] which use the 3*D*-structure of compounds and their conformations (3*D*-position of the compound) [84]. On the other hand, while ML methods have been actively used in the field for the last two decades, new deep learning methods made a huge impact in drug discovery process in the last 5 years [88, 52, 82]. Further discussion of state-of-the-art ML/DL methods are given in Section 6 where we compare our models and benchmark against them.

### 2.2 Topological Data Analysis

TDA and tools of persistent homology (PH) have recently emerged as powerful approaches for ML, allowing us to extract complementary information on the observed objects, especially, from graph-structured data. In particular, PH has become popular for various ML tasks such as clustering, classification, and anomaly detection, with a wide range of applications including material science [68, 43], insurance [99, 46], finance [55], and cryptocurrency analytics [33, 4, 73]. (For more details see surveys [6, 22] and TDA applications library [34]) Furthermore, it has become a highly active research area to integrate PH methods into geometric deep learning (GDL) in recent years [41, 100, 19, 23]. Most recently, the emerging concepts of *multipersistence* (MP) are proposed to advance the success of single parameter persistence (SP) by allowing the use of more than one domain function in the process to produce more granular topological descriptors of the data. However, the MP theory is not sufficiently mature as it suffers from the nonexistence of the barcode decomposition relating to the partially ordered structure of the index set {(*α*_*i*_, *β*_*j*_)} [57, 89]. The existing approaches remedy this issue via slicing technique by studying one-dimensional fibers of the multiparameter domain [18], but choosing these directions suitably and computing restricted SP vectorizations are computationally costly which makes the approach inefficient in real life applications. There are several promising recent studies in this direction [11, 93, 24], but these approaches fail to provide a practical topological summary such as “multipersistence diagram”, and an effective MP vectorization to be used in real life applications.

### 2.3 TDA in Virtual Screening

In [16, 15, 14], the authors obtained successful results by integrating single persistent homology outputs with various ML models. Furthermore, in [50], the authors used multipersistence homology with fibered barcode approach in the 3*D* setting and obtained promising results. In the past few years, TDA tools were also successfully combined with various deep learning models for SBVS and property prediction [71, 72]. In [66, 45, 61, 95, 62], the authors successfully used TDA methods to generate powerful molecular descriptors. Then, by using these descriptors, they highly boosted the performance of various ML/DL models and outperformed the existing models in several benchmark datasets. For a discussion and comparison of TDA techniques with other approaches in virtual screening and property prediction, see the review article [70]. In this paper, we follow a different approach and propose a framework by adapting multipersistence homology to VS process which produces fine topological fingerprints which are highly suitable for ML/DL methods.

## 3 Background

We first provide the necessary TDA background for our machinery. While our techniques are applicable to various forms of data, e.g., point clouds and images (for details, see Section B.2), here we focus on the graph setup in detail with the idea of mapping the atoms and bonds that make up a compound into a set of nodes and edges that represent an undirected graph.

### 3.1 Persistent Homology

Persistent homology (PH) is a key approach in TDA, allowing us to extract the evolution of subtler patterns in the data shape dynamics at multiple resolution scales which are not accessible with more conventional, non-topological methods [17]. In this part, we go over the basics of PH machinery on graph-structured data. For further background on PH, see Appendix A.1 and [27, 30].

For a given graph 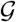, consider a nested sequence of subgraphs 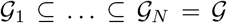. For each 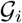, define an abstract simplicial complex 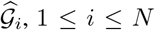, yielding a *filtration*, a nested sequence of simplicial complexes 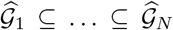. This step is crucial in the process as one can inject domain information to the machinery exactly at this step by using a filtering function from domain, e.g., atomic mass, partial charge, bond type, electron affinity, ionization energy (See Appendix A.1). After getting a filtration, one can systematically keep track of the evolution of topological patterns in the sequence of simplicial complexes 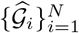. A *k*-dimensional topological feature (or *k*-hole) may represent connected components (0-dimension), loops (1-dimension) and cavities (2-dimension). For each *k*-dimensional topological feature *σ*, PH records its first appearance in the filtration sequence, say 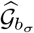, and first disappearence in later complexes, 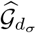 with a unique pair (*b*_*σ*_, *d*_*σ*_), where 1 ≤ *b*_*σ*_ < *d*_*σ*_ ≤ *N*. We call *b*_*σ*_ *the birth time* of *σ* and *d*_*σ*_ *the death time* of *σ*. We call *d*_*σ*_ − *b*_*σ*_ *the life span* (or persistence) of *σ*. PH records all these birth and death times of the topological features in *persistence diagrams*. Let 0 ≤ *k* ≤ *D* where *D* is the highest dimension in the simplicial complex 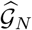. Then *k*^*th*^ persistence diagram 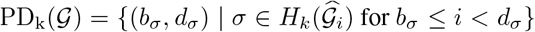. Here, 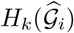 represents the *k^th^ homology group* of 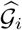 which keeps the information of the *k*-holes in the simplicial complex 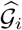. Most common dimensions used in practice are 0 and 1, i.e., 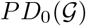 and 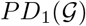. For sake of notations, further we skip the dimension (subscript *k*). With the intuition that the topological features with long life spans (persistent features) describe the hidden shape patterns in the data, these persistence diagrams provide a unique topological fingerprint of 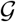. We give the further details of the PH machinery and how to integrate domain information into the process in Appendix A.1.

### 3.2 Multidimensional Persistence

MultiPersistence (MP) significantly boosts the performance of the single parameter persistence technique described in Appendix A.1. The reason for the term “single” is that we are filtering the data in only one direction 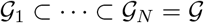. As explained in Appendix A.1, the construction of the filtration is the key step to inject domain information to process and to find the hidden patterns of the data. If one uses a function 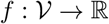 which has valuable domain information, then this induces a single parameter filtration as above. However, various data have more than one domain function to analyze the data, and using them simultaneously would give a much better understanding of the hidden patterns. For example, if we have two functions 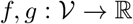 (e.g., atomic mass and partial charge) with valuable complementary information of the network (compound), MP idea is presumed to produce a unique topological fingerprint combining the information from both functions. These pair of functions *f, g* induces a multivariate filtering function 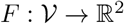 with *F*(*v*) = (*f*(*v*), *g*(*v*)). Again, one can define a set of nondecreasing thresholds 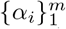 and 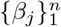 for *f* and *g* respectively. Let 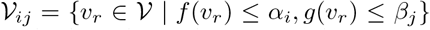, i.e., 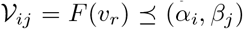. Define 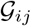 to be the induced subgraph of 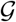 by 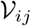, i.e., the smallest subgraph of 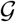 generated by 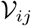. Then, instead of a single filtration of complexes 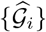, we get a *bifiltration* of complexes 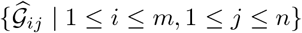 which is a *m* × *n* rectangular grid of simplicial complexes. Again, the MP idea is to keep track of the *k*-dimensional topological features in this grid 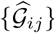 by using the corresponding homology groups 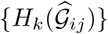 (MP module).

As noted in Section 2, because of the technical problems related to partially ordered structure of the MP module, the MP theory has no sound definition yet (e.g., birth/death time of a topological feature in MP grid), and there is no effective way to facilitate this promising idea in real life applications. In the following, we overcome this problem by producing highly effective fingerprints by utilizing the *slicing* idea in the MP grid in a structured way.

## 4 New Topological Fingerprints of the Compounds with Multipersistence

ToDD framework produces fingerprints of compounds as multidimensional vectors by expanding single persistence (SP) fingerprints (Appendix A.1). While our construction is applicable and suitable for various forms of data, here we focus on graphs, and in particular, compounds for virtual screening. We obtain a 2*D* matrix (or 3D array) for each compound as its fingerprint employing 2 or 3 functions/weights (e.g., atomic mass, partial charge, bond type, electron affinity, ionization energy) to perform graph filtration. We explain how to generalize our framework to other types of data in Appendix B.2. In Appendix B.4, we construct the explicit examples of MP Fingerprints for most popular SP Vectorizations, e.g., Betti, Silhouette, Landscapes.

Our framework basically expands a given SP vectorization to a multidimensional vector by utilizing MP approach. In technical terms, by using the existing SP vectorizations, we produce multidimensional vectors by effectively using one of the filtering direction as *slicing direction* in the multipersistence module. We explain our process in three steps.

*Step 1 - Bifiltrations*: This step basically corresponds to obtaining relevant *substructures* from the given compound in an organized way. Here, we give the computationally most feasible method, called *sublevel bifiltration* with 2 functions. Depending on the task and dataset, the other filtration types or more functions/weights can be more useful. In Section B.5, we give details for other filtration methods we use in our experiments. i.e., Vietoris-Rips (distance) and weight filtration.

Let 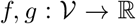 be two filtering functions with threshold sets 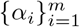 and 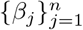 respectively (e.g., *f* is atomic mass, and *g* is partial charge). Let 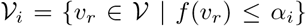 and let 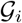 be the induced subgraph of 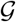 by 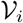, i.e. add any edge in 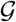 whose endpoints are in 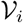. Similarly, let 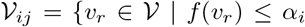 and 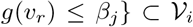. Let 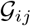 be the induced subgraph of 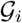 by 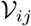. Then, define 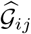 as *the clique complex* of 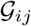 (See Section A.1). In particular, by using the first function (*f*), we filter 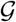 in one (say vertical) direction 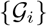. Then, by using the second function (*g*), we filter each 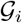 in horizontal direction and obtain a bifiltration 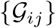. These subgraphs 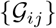 represent the induced substructures of the compound 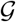 by using the filtering functions *f* and *g*.

In Figure 2 and 3, we give an example of sublevel bifiltration of the compound cytosine by atomic number and partial charge functions. In Figure 2, atom types are coded by their color. Atomic numbers are given in the parenthesis. White=Hydrogen (1), Gray=Carbon (6), Blue=Nitrogen (7), and Red=Oxygen (8). The decimal numbers next to atoms represent their partial charges.

**Figure 2:**
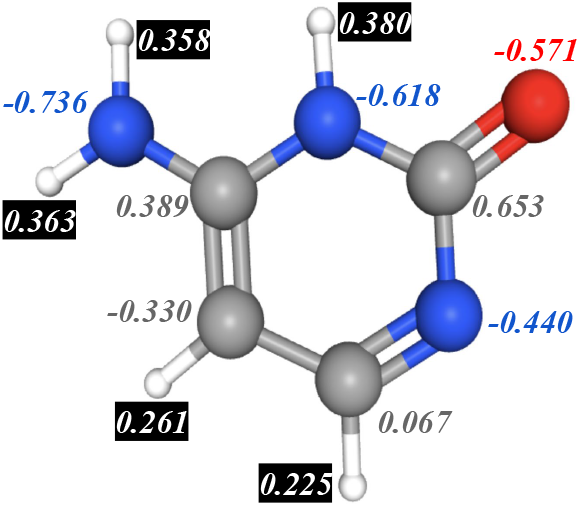
Cytosine. Atom types are coded by their color: White=Hydrogen, Gray=Carbon, Blue=Nitrogen, and Red=Oxygen. The decimal numbers next to atoms represent their partial charges.

**Figure 3:**
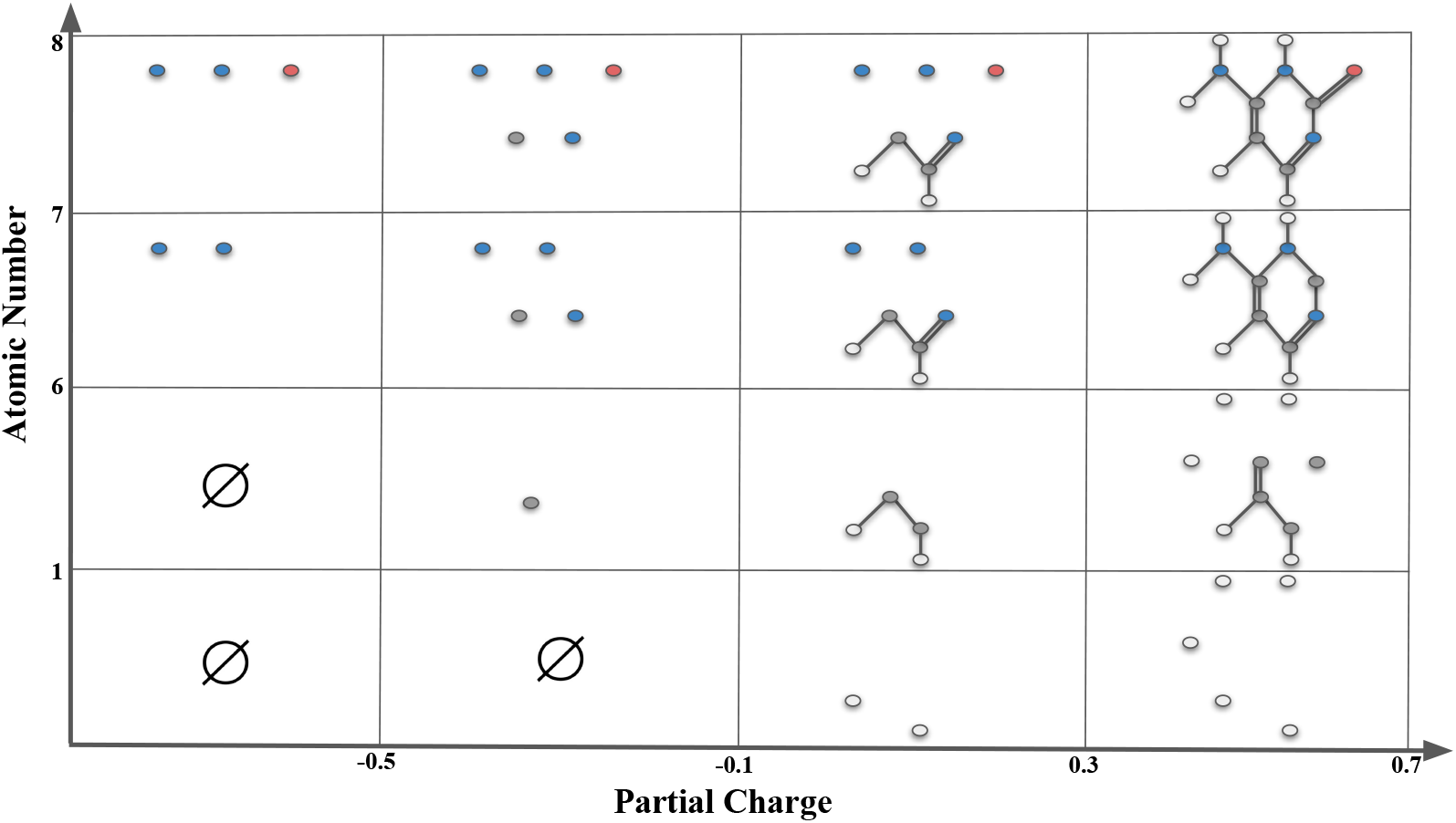
**Sublevel bifiltration of cytosine** is induced by filtering functions atomic charge *f* and atomic number *g*. In the horizontal direction, thresholds *α* = −0.5, −0.1, +0.3, +0.7 filters the compound into substructures *f*(*v*) ≤ *α* with respect to their partial charge. In the vertical direction, thresholds *β* = 1, 6, 7, 8 filters the compound in the substructures *g*(*v*) ≤ *β* with respect to atomic numbers. Each box Δ_*α*,*β*_ indexed by their upper right coordinates (*α*, *β*) representing the substructure Γ_*α*,*β*_ = {*f*(*v*) ≤ *α, g*(*v*) ≤ *β*}. Whenever two nodes (atoms) are in the substructure, if there is an edge (bond) between them in the original compound, we include the edge in the substructure.

*Step 2 - Persistence Diagrams*: After constructing the bifiltration 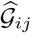, the second step is to obtain persistence diagrams for each row. By restricting the bifiltration to a single row, for each 1 ≤ *i*_0_ ≤ *m*, one obtains a single filtration 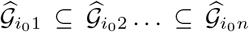 in horizontal direction. This is called a *horizontal slice* in the bipersistence module. Each such single filtration induces a persistence diagram 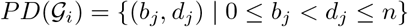. This produces *m* persistence diagrams 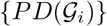. Notice that one can consider 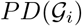 as the single persistence diagram of the “substructure” 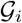 filtered by the second function *g* (See Section A.1).

*Step 3 - Vectorization*: The final step is to use a vectorization on these *m* persistence diagrams. Let *φ* be a single persistence vectorization, e.g., Betti, Silhouette, Entropy, Persistence Landscape or Persistence Image. Specifically, we use Betti to ease computational complexity. By applying the chosen SP vectorization *φ* to each PD, we obtain a function 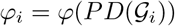 where in most cases it is a single variable function on the threshold domain [0, *n*], i.e., *φ*_*i*_ : [1, *n*] → ℝ. The number of thresholds *m, n* are important as it determines the size of our topological fingerprint. As most such vectorizations are induced from a discrete set of points 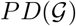, it is common to express them as vector in the form 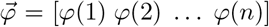. In the examples in Section B.4, we explain this conversion explicitly for different vectorizations. Hence, we obtain a vector 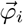 of size 1 × *n* for each row 1 ≤ *i* ≤ *m*.

Now, we can define our topological fingerprint **M**_*φ*_ which is a 2*D*-vector (a matrix)

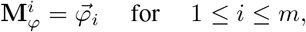

where 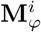 is the *i*^*th*^-row of **M**_*φ*_. Hence, **M**_*φ*_ is a 2*D*-vector of size *m*×*n*. Each row 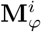 is the vectorization of the persistence diagram 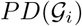 via the SP vectorization method *φ*. We use the first filtering function *f* to get a finer look at the graph as it defines the subgraphs 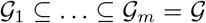. Then, by using the second function *g* on each 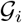, we record the evolution of topological features in each 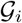 as 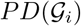. While this construction gives our 2*D* (matrix) fingerprints **M**_*φ*_, one can also use 3 functions/weights for filtration and obtain a finer 3*D* (array) topological fingerprint (Section B.3).

In a way, we look at 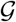 with a 2*D* resolution (functions *f* and *g* as lenses) and keep track of the evolution of topological features in the induced substructures 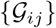. The main advantage of this technique is that the outputs are fixed size multidimensional vectors for each dataset which are suitable for various ML/DL models.

### 4.1 Stability of the MP Fingerprints

We further show that when the source single parameter vectorization *φ* is stable, then so is its induced MP Fingerprint **M**_*φ*_. (We give the details of stability notion in persistence theory and proof of the following theorem in Section B.1.)

#### Theorem

*Let φ be a stable SP vectorization. Then, the induced MP Fingerprint* **M**_*φ*_ *is also stable, i.e., with the notation introduced in Section B.1, there exists* 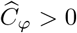 *such that for any pair of graphs* 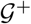 *and* 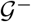, *we have the following inequality*.

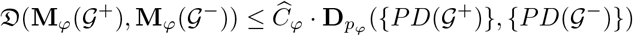

## 5 Datasets

### Cleves-Jain

This is a relatively small dataset [26] that has 1149 compounds.^*^ There are 22 different drug targets, and for each one of them the dataset provides only 2-3 template active compounds dedicated for model training, which presents a few-shot learning task. All targets {*q*} are associated with 4 to 30 active compounds {*L_q_*} dedicated for model testing. Additionally, the dataset contains 850 decoy compounds (*D*). The aim is for each target *q*, by using the templates, to find the actives *L_q_* among the pool combined with decoys *L_q_* ∪ *D*, i.e., same decoy set *D* is used for all targets.

### DUD-E Diverse

DUD-E (Directory of Useful Decoys, Enhanced) dataset [67] is a comprehensive ligand dataset with 102 targets and approximately 1.5 million compounds.^*^ The targets are categorized into 7 classes with respect to their protein type. The “Diverse subset” of DUD-E contains targets from each category to give a balanced benchmark dataset for VS methods. Diverse subset contains 116,105 compounds from 8 target and 8 decoy sets. One decoy set is used per target.

More detailed information about each dataset can be found in Appendix C.1.

## 6 Experiments

### 6.1 Setup

#### Macro Design

We construct different ToDD (Topological Drug Discovery) models, namely ToDD-ViT, ToDD-ConvNeXt and ToDD-RF to test the generalizability and scalability of topological features while employing different ML models and training datasets of various sizes. Many neural network architectural choices and ML models can be incorporated in our ToDD method. ToDD-ViT and ToDD-ConvNeXt are Triplet network architectures with Vision Transformer (ViT_b_16) [28] and ConvNeXt_tiny models [63], pretrained on ILSVRC-2012 ImageNet, serving as the backbone of the Triplet network. MP signatures of compounds are applied nearest neighbour interpolation to increase their resolutions to 224^2^, followed by normalization. We only use GaussianBlur with kernel size 5^2^ and standard deviation 0.05 as a data augmentation technique. Transfer learning via fine-tuning ViT_b_16 and ConvNeXt_tiny models using Adam optimizer with a learning rate of 5e-4, no warmup or layerwise learning rate decay, cosine annealing schedule for 5 epochs, stochastic weight averaging for 5 epochs, weight decay of 1e-4, and a batch size of 64 for 10 epochs in total led to significantly better performance in Enrichment Factor and ROC-AUC scores compared to training from scratch. The performance of all models was assessed by 5-fold cross-validation (CV).

Due to structural isomerism, molecules with identical molecular formulae can have the same bonds, but the relative positions of the atoms differ [76]. ViT has much less inductive bias than CNNs, because locality and translation equivariance are embedded into each layer throughout the entire network in CNNs, whereas in ViT self-attention layers are global and only MLP layers are translationally equivariant and local [28]. Hence, ViT is more robust to distinct arrangements of atoms in space, also referred to as molecular conformation. On a small-scale dataset like Cleves-Jain, ViT exhibits impressive performance. However, the memory and computational costs of dot-product attention blocks of ViT grow quadratically with respect to the size of input, which limits its application on large-scale datasets [60, 83]. Another major caveat is that the number of triplets grows cubically with the size of the dataset. Since ConvNeXt depends on a fully-convolutional paradigm, its inherently efficient design is viable on large-scale datasets like DUD-E Diverse. As depicted in Figure 4, ToDD-ViT and ToDD-ConvNeXt project semantically similar MP signatures of compounds from data manifold onto metrically close embeddings using triplet margin loss with margin *α* = 1.0 and norm *p* = 2 as provided in Equation 1. Analogously, semantically different MP signatures are projected onto metrically distant embeddings.

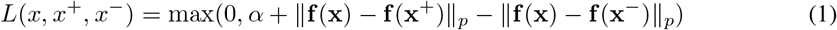

**Figure 4:**
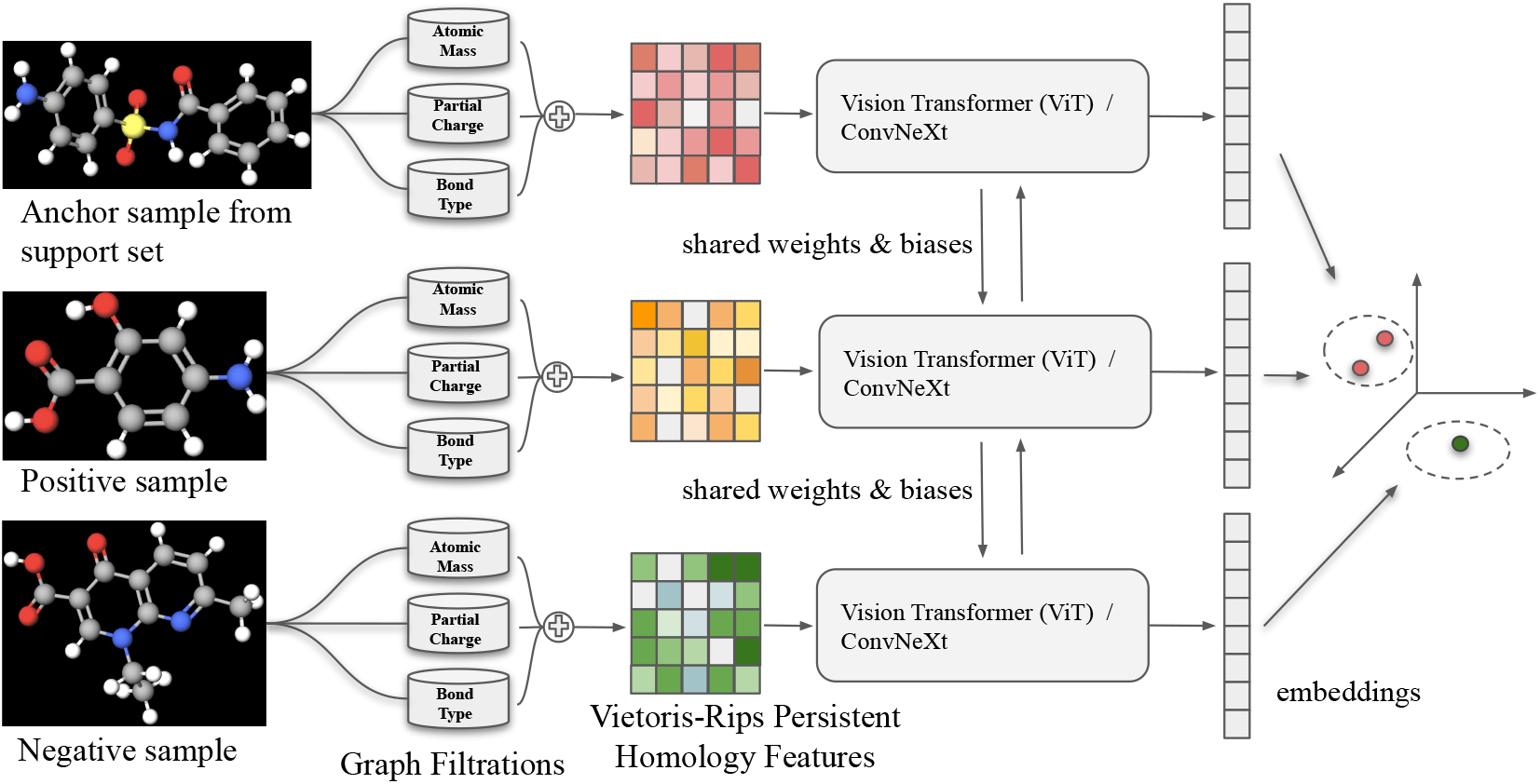
End-to-end model pipeline. Anchor sample, *x*, and positive sample, *x*^+^, are compounds that can bind to the same drug target, whereas negative sample, *x*^−^, is a decoy. 2*D* graph representation of each compound is decomposed into subgraphs induced by the periodic properties: atomic mass, partial charge and bond type. Potentially these domain functions can be augmented using other periodic properties such as ionization energy and electron affinity as well as using molecular information such as chirality, orbital hybridization, number of Hydrogen bonds or number of conjugated bonds at the cost of computational complexity. Subgraphs may have isolated nodes and edges. Our MP framework establishes Vietoris-Rips complexes for each subgraph and provides MP signatures (topological fingerprints) of the compounds. Both ToDD-ViT and ToDD-ConvNeXt can encode the pair of distances between a positive query and a negative query against an anchor sample from the support set.

#### Sampling Strategy

Learning metric embeddings via triplet margin loss on large-scale datasets poses a special challenge in sampling all distinct triplets (*x*, *x*^+^, *x*^−^), and collecting them into a single database causes excessive overhead in computation time and memory. Let *P* be a set of compounds, *x*_*i*_ denotes a compound that inhibits the drug target *i*, and *d*_*ij*_ = *d*(*x*_*i*_, *x*_*j*_) ∈ ℝ denotes a pairwise distance measure which estimates how strongly *x*_*i*_ ∈ *P* is similar to *x*_*j*_ ∈ *P*. The distance metric can be chosen as Euclidean distance, cosine similarity or dot-product between embedding vectors. We use pairwise Euclidean distance computed by the pretrained networks in the implementation. Since triplets (*x*, *x*^+^, *x*^−^) with *d*(*x*, *x*^−^) > *d*(*x*, *x*^+^) + *α* have already negative queries sufficiently distant to the anchor compounds from the support set in the embedding space, they are not sampled to create the training dataset. We only sample triplets that satisfy *d*(*x*, *x*^−^) < *d*(*x*, *x*^+^) (where negative query is closer to the anchor than the positive) and *d*(*x*, *x*^+^) < *d*(*x*, *x*^−^) < *d*(*x*, *x*^+^) + *α* (where negative query is more distant to the anchor than the positive, but the distance is less than the margin).

#### Enrichment Factor

(EF) is the most common performance evaluation metric for VS methods [90]. VS method *φ* ranks compounds in the database by their similarity scores. We measure the similarity score using the inverse of Euclidean distance between the embeddings of an anchor and drug candidate. Let *N* be the total number of ligands in the dataset, *A*_*φ*_ be the number of true positives (i.e., correctly predicted active ligands) ranked among the top *α*% of all ligands (*N*_*α*_ = *N* · *α*%) and *N*_actives_ be the number of active ligands in the whole dataset. Then, 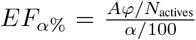. In other words, *EF*_*α*%_ interprets as how much VS method *φ enrich* the possibility of finding active ligand in the first *α*% of all ligands with respect to the random guess. This method is also known as *precision at k* in the literature. With this definition, the max score for *EF*_*α*%_ is 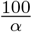, i.e., 100 for *EF*_1%_ and 20 for *EF*_5%_.

### 6.2 Experimental Results

We compare our methods against the 23 state-of-the-art baselines (see Appendix C.2).

Relative gains are relative to the next best performing model. Based on the results (mean and standard deviation of EF scores evaluated by CV) reported in Table 1 and 2, we observe the following:

- ToDD models consistently achieve the best performance on both Cleves-Jain and DUD-E Diverse datasets across all targets and *EF*_*α*%_ levels.
- ToDD learns hierarchical topological representations of compounds using their atoms’ periodic properties, and captures the complex chemical properties essential for high-throughput VS. These strong hierarchical topological representations enable ToDD to become a model-agnostic method that is extensible to state-of-the-art neural networks as well as ensemble methods like random forests (RF).
- For small-scale datasets such as Cleves-Jain, RF is less accurate than ViT despite regularization by bootstrapping and using pruned, shallow trees, because small variations in the data may generate significantly different decision trees. For large-scale datasets such as DUD-E Diverse, ToDD-RF and ToDD-ConvNeXt exhibit comparable performances except for: CP3A4, GCR and HIVRT. We conclude that transformer-based models are more robust than convolutional models and RF classifiers despite increased computation time.

### 6.3 Computational Complexity

Computational complexity (CC) of MP Fingerprint 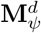 depends on the vectorization] *Ψ* used and the number *d* of the filtering functions one uses. CC for a single persistence diagram *PD*_*k*_ is 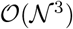 [74], where 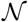 is the number of *k*-simplices. If *r* is the resolution size of the multipersistence grid, then 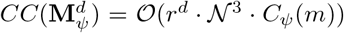 where *C*_*Ψ*_(*m*) is CC for *Ψ* and *m* is the number of barcodes in *PD*_*k*_, e.g., if *Ψ* is Persistence Landscape, then *C*_*Ψ*_(*m*) = *m*^2^ [13] and hence CC for MP Landscape with three filtering functions (*d* = 3) is 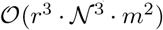. On the other hand, for MP Betti summaries, one does not need to compute persistence diagrams, but the rank of homology groups in the MP module. Hence, for MP Betti summary, the computational complexity is indeed much lower by using minimal representations [58, 51]. To expedite the execution time, the feature extraction task is distributed across the 8 cores of an Intel Core i7 CPU (100GB RAM) running in a multiprocessing process. See Appendix C.4 for an additional analysis of computation time to extract MP fingerprints from the datasets. Furthermore, all ToDD models require substantially fewer computational resources during training compared to current graph-based models that encode a compound through mining common molecular fragments, a.k.a., motifs [47]. Training time of ToDD-ViT and ToDD-ConvNeXt for each individual drug target takes less than 1 hour on a single GPU (*N*VIDIA RTX 2080 Ti).

### 6.4 Ablation Study

We tested a number of ablations of our model to analyze the effect of its individual components and to further investigate the effectiveness of our topological fingerprints.

#### Multimodal Learning

We first address the question of how adding different domain information improves the model performance. In Appendix C.3, we demonstrate one-by-one the importance of each modality (atomic mass, partial charge and bond type) used for graph filtration to the classification of each target. We find that their importance varies across targets in a unimodal setting, but the orthogonality of these information sources offers significant gain in EF scores when the MP signatures learned from each modality are integrated into a joined multimodal representation. Tables 5, 6, 7 and 8 provide detailed results for the performance of each modality across all drug targets.

**Table 5:** EF 2% values and ROC-AUC scores across different modalities on Cleves-Jain dataset using **ToDD-RF**.

| Target | Atomic Mass | Partial Charge | Bond Type | Atomic Mass & Partial Charge | All Modalities |
| --- | --- | --- | --- | --- | --- |
| a | 33.3 | 33.3 | 33.3 | 33.3 | 41.7 |
| b | 25.0 | 29.5 | 31.8 | 27.3 | 25.0 |
| c | 19.2 | 7.7 | 15.4 | 26.9 | 34.6 |
| d | 33.3 | 33.3 | 41.7 | 50.0 | 50.0 |
| e | 30.0 | 30.0 | 30.0 | 40.0 | 40.0 |
| f | 25.0 | 50.0 | 37.5 | 50.0 | 37.5 |
| g | 30.0 | 30.0 | 30.0 | 30.0 | 40.0 |
| h | 40.0 | 50.0 | 30.0 | 50.0 | 50.0 |
| i | 40.0 | 40.0 | 30.0 | 40.0 | 40.0 |
| j | 17.9 | 39.3 | 35.7 | 28.6 | 28.6 |
| k | 21.4 | 21.4 | 17.9 | 35.7 | 32.1 |
| l | 15.0 | 15.0 | 15.0 | 30.0 | 25.0 |
| m | 44.4 | 50.0 | 33.3 | 50.0 | 38.9 |
| n | 15.0 | 25.0 | 10.0 | 25.0 | 10.0 |
| o | 21.7 | 20.0 | 25.0 | 23.3 | 23.3 |
| p | 10.9 | 8.7 | 13.0 | 17.4 | 26.1 |
| q | 45.5 | 27.3 | 22.7 | 40.9 | 40.9 |
| r | 42.9 | 42.9 | 42.9 | 39.3 | 32.1 |
| s | 26.7 | 16.7 | 20.0 | 20.0 | 30.0 |
| t | 30.0 | 50.0 | 50.0 | 50.0 | 50.0 |
| u | 33.3 | 38.9 | 27.8 | 38.9 | 50.0 |
| v | 21.4 | 28.6 | 28.6 | 28.6 | 28.6 |
| Mean | 28.3 | 31.3 | 28.3 | 35.2 | 35.2 |
| ROC-AUC | 0.92 | 0.90 | 0.88 | 0.94 | 0.93 |

**Table 6:** EF 2% values and ROC-AUC scores across different modalities on Cleves-Jain dataset using **ToDD-ViT**.

| Target | Atomic Mass | Partial Charge | Bond Type | Atomic Mass & Partial Charge | All Modalities |
| --- | --- | --- | --- | --- | --- |
| a | 25.0 | 33.3 | 33.3 | 33.3 | 50.0 |
| b | 20.5 | 6.8 | 34.1 | 6.8 | 34.1 |
| c | 11.5 | 15.4 | 23.1 | 7.7 | 46.2 |
| d | 25.0 | 41.7 | 50.0 | 33.3 | 50.0 |
| e | 40.0 | 20.0 | 30.0 | 20.0 | 30.0 |
| f | 25.0 | 25.0 | 37.5 | 25.0 | 50.0 |
| g | 20.0 | 30.0 | 40.0 | 30.0 | 50.0 |
| h | 40.0 | 50.0 | 50.0 | 50.0 | 50.0 |
| i | 30.0 | 20.0 | 20.0 | 20.0 | 50.0 |
| j | 10.7 | 14.3 | 21.4 | 17.9 | 21.4 |
| k | 21.4 | 21.4 | 25.0 | 21.4 | 39.3 |
| l | 15.0 | 30.0 | 30.0 | 40.0 | 35.0 |
| m | 22.2 | 44.4 | 22.2 | 44.4 | 50.0 |
| n | 10.0 | 25.0 | 15.0 | 20.0 | 35.0 |
| o | 20.0 | 16.7 | 16.7 | 18.3 | 20.0 |
| p | 26.1 | 17.4 | 26.1 | 15.2 | 32.6 |
| q | 36.4 | 36.4 | 50.0 | 36.4 | 18.2 |
| r | 14.3 | 17.9 | 42.9 | 21.4 | 32.1 |
| s | 30.0 | 13.3 | 23.3 | 13.3 | 33.3 |
| t | 20.0 | 30.0 | 50.0 | 30.0 | 50.0 |
| u | 33.3 | 38.9 | 38.9 | 33.3 | 50.0 |
| v | 21.4 | 28.6 | 28.6 | 21.4 | 42.9 |
| Mean | 23.5 | 26.2 | 32.2 | 25.4 | 39.5 |
| ROC-AUC | 0.87 | 0.85 | 0.86 | 0.84 | 0.90 |

**Table 7:** EF 1% values and ROC-AUC scores across different modalities on DUD-E Diverse using **ToDD-RF**.

| Model | Atomic Mass |  | Partial Charge |  | Bond Type |  | Atomic Mass & Partial Charge |  | All Modalities |  |
| --- | --- | --- | --- | --- | --- | --- | --- | --- | --- | --- |
|  | EF 1% | ROC-AUC | EF 1% | ROC-AUC | EF 1% | ROC-AUC | EF 1% | ROC-AUC | EF 1% | ROC-AUC |
| AMPC | 42.9 | 0.90 | 42.9 | 0.92 | 28.6 | 0.84 | 42.9 | 0.84 | 28.6 | 0.86 |
| CXCR4 | 84.6 | 0.98 | 76.9 | 0.99 | 84.6 | 0.97 | 92.3 | 0.99 | 92.3 | 0.99 |
| KIF11 | 55.0 | 0.96 | 55.0 | 0.98 | 70.0 | 0.97 | 70.0 | 0.98 | 75.0 | 0.98 |
| CP3A4 | 54.1 | 0.96 | 48.6 | 0.87 | 40.5 | 0.93 | 62.2 | 0.95 | 67.6 | 0.96 |
| GCR | 54.4 | 0.96 | 43.9 | 0.95 | 57.9 | 0.97 | 63.2 | 0.97 | 78.9 | 0.97 |
| AKT1 | 62.8 | 0.97 | 86.0 | 0.99 | 79.1 | 0.98 | 81.4 | 0.98 | 90.7 | 0.99 |
| HIVRT | 53.1 | 0.90 | 46.9 | 0.93 | 64.1 | 0.97 | 54.7 | 0.91 | 64.1 | 0.95 |
| HIVPR | 85.0 | 0.99 | 86.4 | 0.99 | 78.6 | 0.99 | 87.9 | 0.99 | 92.1 | 0.99 |
| **Mean** | 61.5 | 0.95 | 60.8 | 0.95 | 62.9 | 0.95 | 69.3 | 0.95 | 73.7 | 0.96 |

**Table 8:** EF 1% values and ROC-AUC scores across different modalities on DUD-E Diverse using **ToDD-ConvNeXt**.

| Model | Atomic Mass |  | Partial Charge |  | Bond Type |  | Atomic Mass & Partial Charge |  | All Modalities |  |
| --- | --- | --- | --- | --- | --- | --- | --- | --- | --- | --- |
|  | EF 1% | ROC-AUC | EF 1% | ROC-AUC | EF 1% | ROC-AUC | EF 1% | ROC-AUC | EF 1% | ROC-AUC |
| AMPC | 30.8 | 0.90 | 15.4 | 0.83 | 30.8 | 0.73 | 38.5 | 0.92 | 46.2 | 0.81 |
| CXCR4 | 52.0 | 0.98 | 44.0 | 0.92 | 32.0 | 0.94 | 60.0 | 0.97 | 84.0 | 0.99 |
| KIF11 | 47.5 | 0.96 | 45.0 | 0.88 | 37.5 | 0.92 | 60.0 | 0.96 | 72.5 | 0.97 |
| CP3A4 | 19.2 | 0.86 | 17.8 | 0.86 | 15.1 | 0.86 | 28.8 | 0.90 | 28.8 | 0.91 |
| GCR | 25.7 | 0.95 | 30.1 | 0.95 | 19.5 | 0.94 | 43.4 | 0.96 | 46.0 | 0.97 |
| AKT1 | 60.0 | 0.91 | 51.8 | 0.91 | 41.2 | 0.96 | 77.6 | 0.99 | 81.2 | 0.98 |
| HIVRT | 26.6 | 0.93 | 21.9 | 0.89 | 17.2 | 0.94 | 35.9 | 0.94 | 37.5 | 0.95 |
| HIVPR | 65.6 | 0.98 | 50.9 | 0.97 | 45.5 | 0.94 | 70.3 | 0.99 | 74.6 | 0.99 |
| **Mean** | 40.9 | 0.93 | 34.6 | 0.90 | 29.9 | 0.90 | 51.8 | 0.95 | 58.8 | 0.95 |

**Table 9:** Clock time performance to extract Vietoris Rips persistent homology features.

| Dataset | Atomic Mass | Partial Charge | Bond Type |
| --- | --- | --- | --- |
| Cleves-Jain | 00 h : 03 min : 21 sec | 00 h : 03 min : 14 sec | 00 h : 01 min : 13 sec |
| DUD-E Diverse | 06 h : 10 min : 51 sec | 05 h : 37 min : 12 sec | 02 h : 14 min : 54 sec |

#### Morgan Fingerprints

We quantitatively analyze the explainability of our models’ success by replacing topological fingerprints computed via multiparameter persistence with the most popular fingerprinting method: Morgan fingerprints. Our results in Appendix C.5 show that ToDD engineers features that represent the underlying attributes of compounds significantly better than the Morgan algorithm to identify the active compounds across all drug targets. We provide detailed tabulated results of our benchmarking study across all drug targets in Tables 10 and 11.

**Table 10:** EF 2% values on Cleves-Jain Dataset using ViT model trained with Morgan fingerprints vs. ToDD fingerprints.

| Target | Morgan | ToDD |
| --- | --- | --- |
| a | 25.0 | 50.0 |
| b | 11.4 | 34.1 |
| c | 3.8 | 46.2 |
| d | 50.0 | 50.0 |
| e | 10.0 | 30.0 |
| f | 37.5 | 50.0 |
| g | 30.0 | 50.0 |
| h | 40.0 | 50.0 |
| i | 20.0 | 50.0 |
| j | 17.9 | 21.4 |
| k | 14.3 | 39.3 |
| l | 45.0 | 35.0 |
| m | 38.9 | 50.0 |
| n | 15.0 | 35.0 |
| o | 13.3 | 20.0 |
| p | 2.2 | 32.6 |
| q | 18.2 | 18.2 |
| r | 14.3 | 32.1 |
| s | 10.0 | 33.3 |
| t | 20.0 | 50.0 |
| u | 27.8 | 50.0 |
| v | 35.7 | 42.9 |
| Mean | 22.7 | 39.5 |
| ROC-AUC | 0.86 | 0.90 |

**Table 11:** EF 1% values and ROC-AUC scores on DUD-E Diverse dataset using ConvNeXt model trained with Morgan fingerprints vs. ToDD fingerprints.

| Model | Morgan |  | ToDD |  |
| --- | --- | --- | --- | --- |
|  | EF 1% | ROC-AUC | EF 1% | ROC-AUC |
| AMPC | 38.5 | 0.87 | 46.2 | 0.81 |
| CXCR4 | 48.0 | 0.97 | 84.0 | 0.99 |
| KIF11 | 57.5 | 0.95 | 72.5 | 0.97 |
| CP3A4 | 20.5 | 0.84 | 28.8 | 0.91 |
| GCR | 46.7 | 0.94 | 46.0 | 0.97 |
| AKT1 | 60.0 | 0.98 | 81.2 | 0.98 |
| HIVRT | 50.0 | 0.96 | 37.5 | 0.95 |
| HIVPR | 61.3 | 0.98 | 74.6 | 0.99 |
| Mean | 47.8 | 0.94 | 58.8 | 0.95 |

#### Network Architecture

We investigated ways to leverage deep metric learning by architecting *i*) a Siamese network trained with contrastive loss, *ii*) a Triplet network trained with triplet margin loss, and *iii*) a Triplet network trained with circle loss. Based on our preliminary experiments, the embeddings learned by *i* and *iii* provide sub-par results for compound classification, hence we use *ii*.

## 7 Conclusion

We have proposed a new idea of the topological fingerprints in VS, allowing for deeper insights into structural organization of chemical compounds. We have evaluated the predictive performance of our ToDD methodology for computer aided drug discovery on benchmark datasets. Moreover, we have demonstrated that our topological descriptors are model-agnostic and have proven to be exceedingly competitive, yielding state-of-the-art results unequivocally over all baselines. A future research direction is to enrich ToDD with different VS modalities, and use it on ultra-large virtual compound libraries. It is important to note that this new way of capturing the chemical information of compounds provides a transformative perspective to every level of the pharmaceutical pipeline from the very early phases of drug discovery to the final stages of formulation in development.

## 8 Acknowledgments

YG, BC, YC and ISD were partially supported by Simons Collaboration Grant # 579977, the National Science Foundation (NSF) under award # ECCS 2039701, the Department of the Navy, Office of Naval Research (ONR) under ONR award # N00014-21-1-2530. Part of YG’s contribution is also based upon work supported by (while serving at) the NSF. Any opinions, findings, and conclusions or recommendations expressed in this material are those of the author(s) and do not necessarily reflect the views of the NSF and/or the ONR.

## Appendix

## A Topological Data Analysis (TDA)

## A.1 Single Parameter Persistent Homology

Here, we give further details on single parameter persistent homology. To sum up, PH machinery is a 3-step process. The first step is the *filtration* step, where one can integrate the domain information to the process. The second step is the *persistence diagrams*, where the machinery records the evolution of topological features (birth/death times) in the filtration, sequence of the simplicial complexes. The final step is the *vectorization* (fingerprinting) where one can convert these records to a function or vector to be used in suitable ML models.

## Constructing Filtrations

As PH is basically the machinery to keep track of the evolution of topological features in a sequence, the most important step is the construction of the sequence 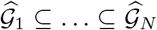. This is the key step where one can inject the valuable domain information to the PH process by using important domain functions (e.g., atomic mass, partial charge). While there are various filtration techniques used for PH machinery on graphs [5, 41], we will focus on two most common methods: *Sublevel/superlevel filtration* and *Vietoris-Rips (VR) filtration*.

For a given unweighted graph (compound) 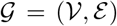 with 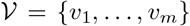 the set of nodes (atoms) and 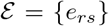 the set of edges (bonds), the most common technique is to use a filtering function 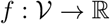 with a choice of thresholds 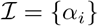 where 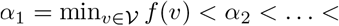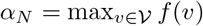. For 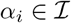, let 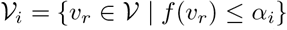 (sublevel sets for *f*). Here, in VS problem, this filtering function *f* can be atomic mass, partial charge, bond type, electron affinity, ionization energy or another important function representing chemical properties of the atoms. One can also use the natural graph induced functions like node degree, betweenness, etc. Let 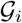 be the induced subgraph of 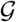 by 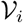, i.e., 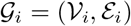 where 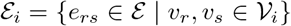. This process yields a nested sequence of subgraphs 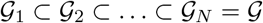. To obtain a filtration, next step is to assign a simplicial complex 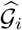 to the subgraph 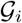. One of the most common techniques is the clique complexes [5]. The clique complex 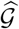 is a simplicial complex obtained from 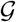 by assigning (filling with) a *k*-simplex to each complete (*k* + 1)-complete subgraph in 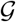, e.g., a 3-clique, a complete 3-subgraph, in 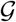 will be filled with a 2-simplex (triangle). This technique is generally known as *sublevel filtration* with clique complexes. As *f*(*v*_*i*_) ≤ *α*_*i*_ condition in the construction gives sublevel filtration, one can similarly use *f*(*v*_*i*_) ≥ *α*_*i*_ condition to define *superlevel filtration*. Similarly, for a weighted graph (bond strength), sublevel filtration on edge weights provides corresponding filtration reflecting the domain information stored in the edge weights [5].

While sublevel/superlevel filtration with clique complexes is computationally cheaper and more common in practise, in this paper, we will essentially use a distance-based filtration technique called *Vietoris-Rips (VR) filtration* where the pairwise distances between the nodes play key role. This technique is computationally more expensive, but gives much finer information about the graph’s intrinsic properties [2]. For a given graph 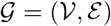, we define the distance between *d*(*v*_*r*_, *v*_*s*_) = *d*_*rs*_ where *d*_*rs*_ is the smallest number of edges required to get from *v*_*r*_ to *v*_*s*_ in 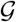. Then, let 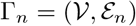 be the graph where 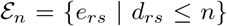, i.e. 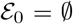 and 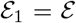 with 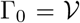 and 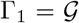. In other words, we start with the nodes of 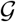, then for any pair of vertices *v*_*r*_, *v*_*s*_ with distance *d*_*rs*_ ≤ n in 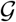, we add an edge *e*_*rs*_ to the graph Γ_*n*_. Then, define the simplicial complex 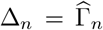, the clique complex of Γ_*n*_. This defines a filtration Δ_0_ ⊂ Δ_1_ ⊂ · · · ⊂ Δ_*K*_ where *K* = max *d*_*rs*_, i.e. the distance between farthest two nodes in the graph 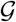. Hence, for *n* ≥ *K*, Δ_*n*_ = Δ_*K*_ which is a (*m* − 1)-simplex as Γ_*K*_ is complete *m*-graph where 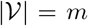. In particular, in this setting, we consider the vertex set 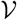 as a point cloud where the distances between the points induced from the graph 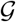. In graph setting, V *R*-filtration is also known as *power filtration* as the graph Γ_*n*_ is also called 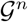, *n*^*th*^ power of 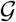.

## Persistence Diagrams

The second step in PH process is to obtain persistence diagrams (PD) for the filtration Δ_0_ ⊂ Δ_1_ ⊂ … ⊂ Δ_*K*_. As explained in Section 3.1, PDs are collection of 2-tuples, marking the birth and death times of the topological features appearing in the filtration, i.e. 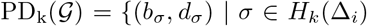 for *b*_*σ*_ ≤ *i* < *d*_*σ*_. This step is pretty standard and there are various software libraries for this task [74].

## Vectorizations (Fingerprinting)

While PH extracts hidden shape patterns from data as persistence diagrams (PD), PDs being collection of points in ℝ^2^ by itself are not very practical for statistical and ML purposes. Instead, the common techniques are by faithfully representing PDs as kernels [54] or vectorizations [39]. One can consider this step as converting PDs into a useful format to be used in ML process as fingerprints of the dataset. This provides a practical way to use the outputs of PH in real life applications. *Single Persistence Vectorizations* transform obtained PH information (PDs) into a function or a feature vector form which are much more suitable for ML tools than PDs. Common single persistence (SP) vectorization methods are Persistence Images [3], Persistence Landscapes [13], Silhouettes [21], Betti Curves and various Persistence Curves [25]. These vectorizations define a single variable or multivariable functions out of PDs, which can be used as fixed size 1*D* or 2*D* vectors in applications, i.e 1 × *n* vectors or *m* × *n* vectors. For example, a Betti curve for a PD with *n* thresholds can also be expressed as 1 × *n* size vectors. Similarly, Persistence Images is an example of 2*D* vectors with the chosen resolution (grid) size. See the examples given in Section B.4 for further details.

## B Multiparameter Persistence (MP) Fingerprints

## B.1 Stability of MP Fingerprints

## Stability of Single Persistence Vectorizations

A given PD vectorization *φ* can be considered as a map from space of persistence diagrams to space of functions, and the stability intuitively represents the continuity of this operator. In other words, stability question is whether a small perturbation in PD cause a big change in the vectorization or not. To make this question meaningful, one needs to define what “small perturbation” means in this context, i.e., a metric in the space of persistence diagrams. The most common such metric is called *Wasserstein distance* (or matching distance) which is defined as follows.

Let 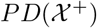 and 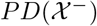 be persistence diagrams two datasets 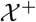 and 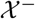 (We omit the dimensions in PDs). Let 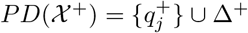 and 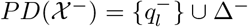 where Δ^±^ represents the diagonal (representing trivial cycles) with infinite multiplicity. Here, 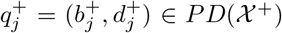 represents the birth and death times of a topological feature *σ*_*j*_ in 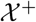. Let 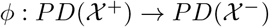 represent a bijection (matching). With the existence of the diagonal Δ^±^ in both sides, we make sure the existence of these bijections even if the cardinalities 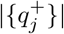 and 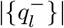 are different.

### Definition B.1

*Let* 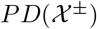 *be persistence diagrams of the datasets* 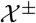, *and* **M** = {*ϕ*} *represent the space of matchings as described above. Then, the p^*th*^ Wasserstein distance* 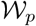 defined as

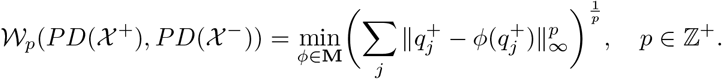

Now, we define stability of vectorizations. A vectorization can be considered as an operator from space of persistence diagrams **P** to space of functions (or vectors) **Y**, e.g., *Ψ* : **P** → **Y**. In particular, when *Ψ* is persistence landscape, 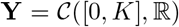 and when *Ψ* is Betti summary, then **Y** = ℝ^*m*^ (See MP Examples in Section B.4) Stability of vectorization *Ψ* basically corresponds to the continuity of *Ψ* as an operator. Let d(., .) be a suitable metric on the space of vectorizations used. Then, we define the stability of *Ψ* as follows:

### Definition B .2

*Let Ψ* : **P** → **Y** *be a vectorization for single persistence diagrams. Let* 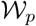 *be the metrics on* **P** *and* **Y** *respectively as described above. Let* 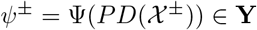. *Then*, *Ψ is called stable if*

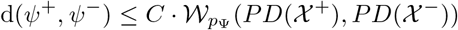

Here, the constant *C* > 0 is independent of 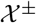. This stability inequality interprets as the changes in the vectorizations are bounded by the changes in PDs. Two nearby persistence diagrams are represented by nearby vectorizations. If a given vectorization *φ* holds such a stability inequality for some *d* and 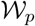, we call *φ* a *stable vectorization* [7]. Persistence Landscapes [13], Persistence Images [3], Stabilized Betti Curves [48] and several Persistence curves [25] are among well-known examples of stable vectorizations.

Now, we are ready to prove the stability of MP Fingerprints given in Section 4.1

Let 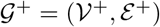 and 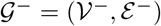 be two graphs. Let *φ* be a stable SP vectorization with the stability equation

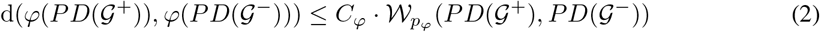

for some 1 ≤ *p*_*φ*_ ≤ ∞. Here, 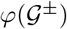 represent the corresponding vectorizations for 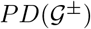 and 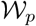 represents Wasserstein-*p* distance as defined in Section B.1.

Now, let 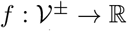 be a filtering function with threshold set 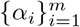. Then, define the sublevel vertex sets 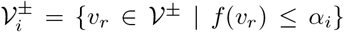. For each 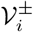, construct the induced VR-filtration 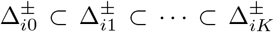 as before. For each 1 ≤ *i*_0_ ≤ *m*, we will have persistence diagram 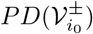 of the filtration 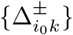.

We define the induced matching distance between the multiple persistence diagrams as

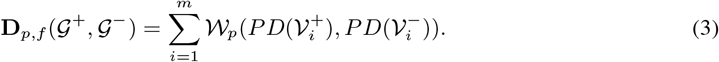

Now, we define the distance between induced MP Fingerprints as

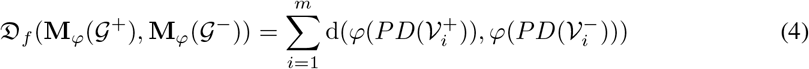

### Theorem B.1

*Let φ be a stable SP vectorization. Then, the induced MP Fingerprint* **M**_*φ*_ *is also stable, i.e., with the notation above, there exists* 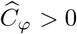 *such that for any pair of graphs* 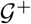 *and* 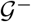, *we have the following inequality*.

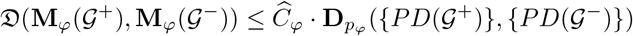

*Proof:* As *φ* is a stable SP vectorization, by Equation 2, for any 1 ≤ *i* ≤ *m*, we have 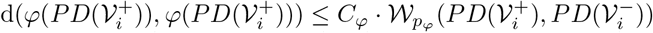 for some *C*_*φ*_ > 0, where 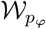 is Wasserstein-*p* distance. Notice that the constant *C*_*φ*_ > 0 is independent of *i*. Hence,

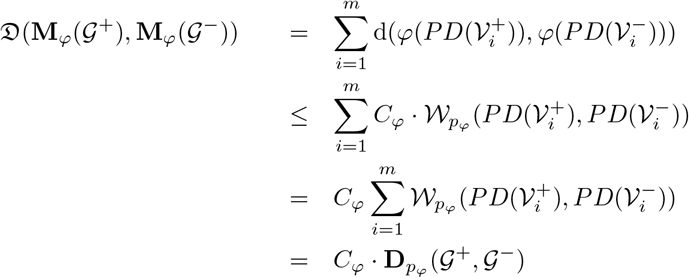

where the first and last equalities are due to Equation 3 and Equation 4, while the inequality follows from Equation 2 which is true for any i. This concludes the proof of the theorem. □

## B.2 MP Fingerprint for Other Types of Data

So far, to keep the exposition focused on VS setting, we described our construction only in the graph setup. However, our framework is suitable for various types of data. Let 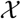 be a an image data or a point cloud. Let 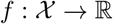 and 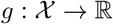 be two filtering functions on 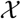. e.g., grayscale function for image data, or density function on point cloud data.

Let 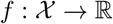 be the filtering function with threshold set 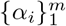.Let 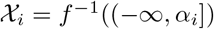. Then, we get a filtering of 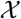 as nested subspaces 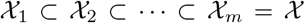. By using the second filtering function, we obtain finer filtrations for each subspace 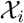 where 1 ≤ *i* ≤ *m*. In particular, fix 1 ≤ *i*_0_ ≤ *m* and let 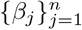 be the threshold set for the second filtering function *g*. Then, by restricting *g* to 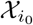, we get a filtering function on 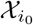, i.e., 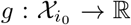 which produces filtering 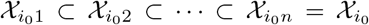. By inducing a simplicial complex 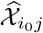 for each 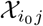, we get a filtration 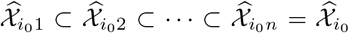. This filtration results in a persistence diagram (PD) 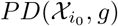. For each 1 ≤ *i* ≤ *m*, we obtain 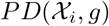. Note that after getting 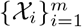 via *f* instead of using second filtering function *g*, one can apply Vietoris-Rips construction based on distance for each 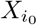 in order to get a different filtration 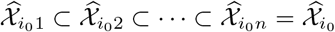.

By using *m* PDs, we follow a similar route to define our MP Fingerprints. Let *φ* be a single persistence vectorization. By applying the chosen SP vectorization *φ* to each PD, we obtain a function 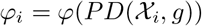 on the threshold domain [*β*_1_, *β*_*n*_], which can be expresses as a 1*D* (or 2*D*) vector in most cases (Section B.4). Let 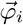 be the corresponding 1 × *k* vector for the function *φ*_*i*_. Define the corresponding MP Fingerprint **M**_*φ*_ as 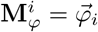 where 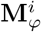 is the *i*^*th*^ row of **M**_*φ*_. In particular, **M**_*φ*_ is a 2*D*-vector (a matrix) of size *m* × *k* where *m* is the number of thresholds for the first filtering function *f*, and *k* is the length of the vector 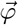.

## B.3 3*D* or higher dimensional MP Fingerprints

If one wants to use two filtering functions and get 3*D*-vectors as the topological fingerprint of the process, the idea is similar. Let 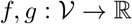 be two filtering functions with threshold sets 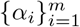 and 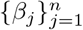 respectively. Let 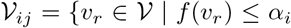 and *g*(*v_r_*) ≤ *β_j_*}. Again, compute all the pairwise distances *d*(*v*_*r*_, *v*_*s*_) = *m*_*rs*_ in 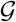 before defining simplicial complexes. Then, for each *i*_0_, *j*_0_, obtain a VR-filtration 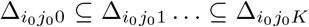 for the vertex set 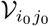 with distances *d*(*v*_*r*_, *v*_*s*_) = *m*_*rs*_ in 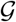. Compute the persistence diagram 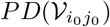 for the filtration 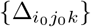. This gives *m* × *n* persistence diagrams 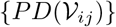. After vectorization, we obtain a 3*D*-vector of size *m* × *n* × *r* as before.

## B.4 Examples of MP Fingerprints

Here, we give explicit constructions of MP Fingerprints for most common SP vectorizations. As noted above, the framework is generalizable and can be applied to most SP vectorization methods. In all the examples below, we use the following setup: Let 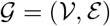 be a graph, and let 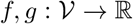 be two filtering functions with threshold sets 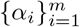 and 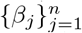 respectively. As explained above, we first apply sublevel filtering with *f* to get a sequence of nested subgraphs, 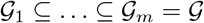. Then, for each 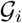, we apply sublevel filtration with *g* to get persistence diagram 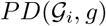. Therefore, we will have *m* PDs. In the examples below, for a given SP vectorization *φ*, we explain how to obtain a vector 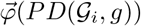, and define the corresponding MP Fingerprint **M**_*φ*_. Note that we skip the homology dimension (subscript *k* for 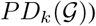 in the discussion. In particular, for each dimension *k* = 0, 1,. . ., we will have one MP Fingerprint 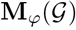 (a matrix) corresponding to 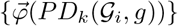. The most common dimensions are *k* = 0 and *k* = 1 in applications.

## MP Landscapes

Persistence Landscapes λ are one of the most common SP vectorizations introduced by [13]. For a given persistence diagram 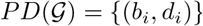, λ produces a function 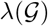 by using generating functions Λ_*i*_ for each 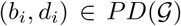, i.e., Λ_*i*_ : [*b*_*i*_, *d*_*i*_] → ℝ is a piecewise linear function obtained by two line segments starting from (*b*_*i*_, 0) and (*d*_*i*_, 0) connecting to the same point 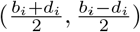. Then, the *Persistence Landscape* function 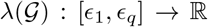 for *t* ∈ [*ϵ*_1_, *ϵ*_*q*_] is defined as

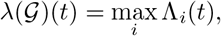

where 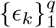 are thresholds for the filtration used.

Considering the piecewise linear structure of the function, 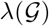 is completely determined by its values at 2*q* − 1 points, i.e., 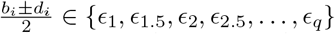 where *ϵ*_*k*.5_ = (*ϵ*_*k*_ + *ϵ*_*k*+1_)/2. Hence, a vector of size 1 × (2*q* − 1) whose entries the values of this function would suffice to capture all the information needed, i.e. 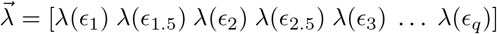

Considering we have threshold set 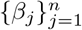 for the second filtering function *g*, 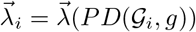 will be a vector of size 1 × 2*n* − 1. Then, as 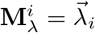 for each 1 ≤ *i* ≤ *m*, MP Landscape 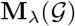 would be a 2*D*-vector (matrix) of size *m* × (2*n* − 1).

## MP Persistence Images

Next SP vectorization in our list is Persistence Images [3]. Different than the most SP vectorizations, Persistence Images produces 2*D*-vectors. The idea is to capture the location of the points in the persistence diagrams with a multivariable function by using the 2*D* Gaussian functions centered at these points. For 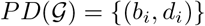, let *ϕ*_*i*_ represent a 2*D*-Gaussian centered at the point (*b_i_, d_i_*) ∈ ℝ^2^. Then, one defines a multivariable function, *Persistence Surface*, 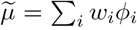 where *w_i_* is the weight, mostly a function of the life span *d*_*i*_ − *b_i_*. To represent this multivariable function as a 2*D*-vector, one defines a *k* × *l* grid (resolution size) on the domain of 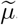 i.e., threshold domain of 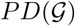 Then, one obtains the *Persistence Image*, a 2*D*-vector (matrix) 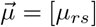 of size *k*×*l* where 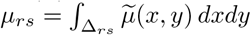 and Δ_*rs*_ is in the is the corresponding pixel (rectangle) *k* × *l* grid.

This time, the resolution size *k* × *l* is independent of the number of thresholds used in the filtering, the choice of *k* and *l* is completely up to the user. Recall that by applying the first function *f*, we have the nested subgraphs 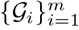. For each 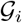, the persistence diagram 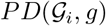 obtained by sublevel filtration with *g* induces a 2*D* vector 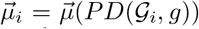 of size *k* × *l*. Then, define MP Persistence Image as 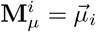, where 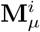 is the *i*^*th*^-floor of the matrix **M**_*μ*_. Hence, 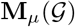 would be a 3*D*-vector of size *m* × *k* × l where *m* is the number of thresholds for the first function *f* and *k* × *l* is the chosen resolution size for the Persistence Image 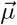.

## MP Betti Summaries

Next, we give an important family of SP vectorizations, Persistence Curves [25]. This is an umbrella term for several different SP vectorizations, i.e., Betti Curves, Life Entropy, Landscapes, et al. Our MP Fingerprint framework naturally adapts to all Persistence Curves to produce multidimensional vectorizations. As Persistence Curves produce a single variable function in general, they all can be represented as 1*D*-vectors by choosing a suitable mesh size depending on the number of thresholds used. Here, we describe one of the most common Persistence Curves in detail, i.e., Betti Curves. It is straightforward to generalize the construction to other Persistence Curves.

Betti curves are one of the simplest SP vectorization as it gives the count of topological feature at a given threshold interval. In particular, *β*_*k*_(Δ) is the total count of *k*-dimensional topological feature in the simplicial complex Δ, i.e., *β*_*k*_(Δ) = *rank*(*H*_*k*_(Δ)). Then, 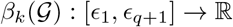 is a step function defined as

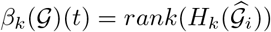

for *t* ∈ [*ϵ*_*i*_, *ϵ*_*i*+1_), where 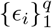 represents the thresholds for the filtration used. Considering this is a step function where the function is constant for each interval [*ϵ*_*i*_, *ϵ*_*i*+1_), it can be perfectly represented by a vector of size 1 × *q* as 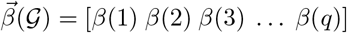.

Then, with the threshold set 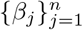 for the second filtering function *g*, 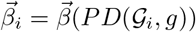 will be a vector of size 1 × *n*. Then, as 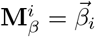 for each 1 ≤ *i* ≤ *m*, MP Betti Summary 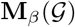 would be a 2*D*-vector (matrix) of size *m* × *n*. In particular, each entry **M**_*β*_ = [*m*_*ij*_] is just the Betti number of the corresponding clique complex in the bifiltration 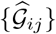, i.e., 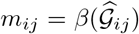. This matrix **M**_*β*_ is also called *bigraded Betti numbers* in the literature, and computationally much faster than other vectorizations [58, 51].

## B.5 MP Vectorization with Other Filtrations

In our paper, other than the simple bifiltration explained in Section 4, we also used the following two filtrations. In the Vietoris-Rips filtration, we use graph geodesic (VR-filtration) as our natural slicing direction. The motivation for this choice is that *VR-filtration captures fine intrinsic structure of the graph by using the pairwise distances between the nodes (atoms)*. With the weight filtration, we can utilize the bond strength of compounds effectively in our construction.

## Vietoris-Rips filtration

Here, we describe our VR construction for 2*D* multipersistence. The construction can easily be extended to 3*D* or higher dimensions (See Appendix B.3). Let 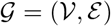 be a graph, and let 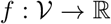 be a filtering function (e.g., atomic mass, partial charge, bond type, electron affinity, ionization energy) with threshold sets 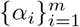. Let 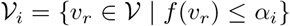. This defines a hierarchy 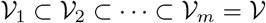 among the nodes with respect to the function *f*.

Before constructing simplicial complexes, compute the distances between each node in graph 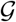, i.e., *d*(*v*_*r*_, *v*_*s*_) = *d*_*rs*_ is the length of the shortest path from *v*_*r*_ to *v*_*s*_ where each edge has length 1. Let *K* = *max d*_*rs*_. Then, for each 1 ≤ *i*_0_ ≤ *m*, define VR-filtration for the vertex set 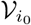 with the distances *d*(*v*_*r*_, *v*_*s*_) = *d*_*rs*_ as described in Section 3.1, i.e., 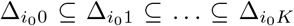 (See Figure 5). This gives *m* × (*K* + 1) simplicial complexes {Δ_*ij*_} where 1 ≤ *i* ≤ *m* and 0 ≤ *j* ≤ *K*.

**Figure 5:**
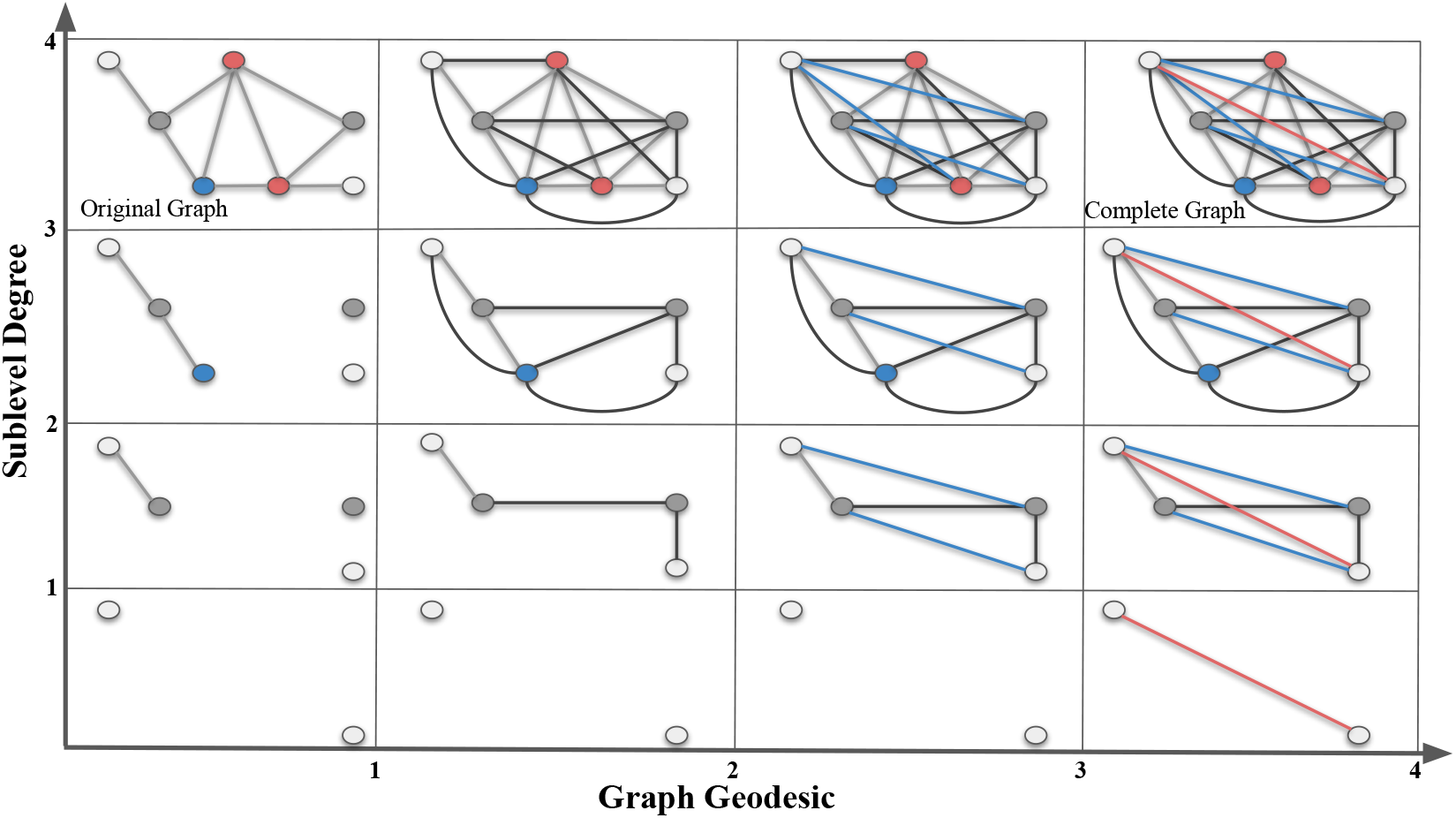
Vietoris-Rips Filtrations. In this toy example, we give a bifiltration composed of a sublevel (vertical) and a VR filtration (horizontal) of a simple graph 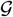 (top box in the first column). In the vertical direction, we apply sublevel filtration by degree function with thresholds 1, 2, 3 and 4. In the horizontal direction, we apply VR-filtration with respect to graph distance (geodesic length). In the first column, we have an (gray) edge between two nodes if their graph distance is 1. In the second column, we have an (black) edge between two nodes if their graph distance is ≤ 2. Blue edges in the third column represent the edges for graph distance 3. Red edges in the last column represent the edges for graph distance 4.

This is called the *bipersistence module*. One can imagine increasing sequence of 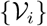 as vertical direction, and induced VR-complexes {Δ_*ij*_} as the horizontal direction. In our construction, we fix the slicing direction as the horizontal direction (VR-direction) in the bipersistence module, and obtain the persistence diagrams in these slices.

In the toy example in Figure 5, we use a small graph 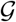 instead of a real compound to keep the exposition simple. Our sublevel filtration (vertical direction) comes from the degree function. Degree of a node is the number of edges incident to it. In the first column, we simply see the single sublevel filtration of 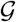 by the degree function. In each row, we develop VR-filtration of the subgraph by using the graph distances between the nodes. Here, graph distance between nodes means the length of the shortest path (geodesic) in the graph where each edge is taken as length 1. Then, in the second column, we add the edges for the nodes whose graph distance is equal to 2. In the third column, we add the (blue) edges for the nodes whose graph distance is equal to 3. Finally, in the last column, we add the (red) edges for the nodes whose graph distance is equal to 4. By construction, all the graphs in the last column must be a complete graph as there is no more edge to add.

After getting the bifiltration, the following steps are the same as in Section 4. In particular, for each 1 ≤ *i*_0_ ≤ *m*, one obtains a single filtration 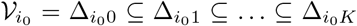 in horizontal direction. This single filtration gives a persistence diagram 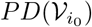 as before. Hence, one obtains *m* persistence diagrams 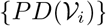. Again, by applying a vectorization *φ*, one obtains *m* row vectors of fixed size *r*, i.e. 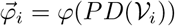. This induces a 2*D*-vector **M**_*φ*_ (a matrix) of size *m* × (*K* + 1) as before.

While computationally more expensive than others, VR-filtration can be very effective for some VS tasks, as it detects the graph distances between atoms, and size of the topological features [59, 2]. Note that VR-filtration when used for unweighted graphs with graph distance is known as “power filtration” in the literature. For further details on VR-filtration, see [5, 27].

## Weight filtration

For a given weighted graph 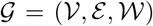, it is common to use edge weights 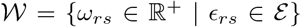 to describe filtration. For example, in our case, one can take bond strength in the compounds as edge weights. By choosing the threshold set similarly 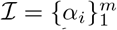 with 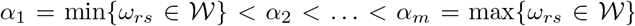. For 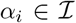, let 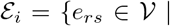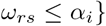. Let 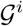 be a subgraph of 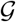 induced by 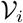. This induces a nested sequence of subgraphs 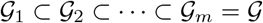.

In the case of weighted graphs, one can apply the MP vectorization framework just by replacing the first filtering (via *f*) with weight filtering. In particular, let 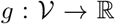 be a filtering function with threshold set 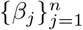. Then, one can first apply weight filtering to get 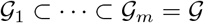 as above, and then apply *f* to each 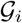 to get a bilfiltration 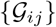 (*m* × *n* resolution). One gets *m* PDs as 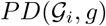 and induce the corresponding **M**_*φ*_. Alternatively, one can change the order by applying *g* first, and get a different filtering 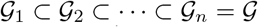 induced by *g*. Then, apply to edge weight filtration to any 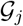, one gets a bifiltration 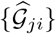 (*n* × *m* resolution) this time. As a result, one gets *n* PDs as 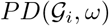 and induce the corresponding **M**_*φ*_. The difference is that in the first case (first apply weights, then *g*), the filtering function plays more important role as **M**_*φ*_ uses 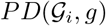 while in the second case (first apply *g*, then apply weights) weights have more important role as **M**_*φ*_ is induced by 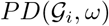. Note also that there is a different filtration method for weighted graphs by applying the following the following VR-complexes method.

In our applications, we used weight filtration to express bond strength in the compounds. Single bond has weight 1, double bond has weight 2, triple bond has weight 3, and finally aromatic bond has weight 4 on the edges.

## C Further Experimental Results

## C.1 Dataset Statistics

## C.2 Baselines

We compare our methods against the 23 state-of-the-art baselines including USR [8], ROCS [38], PS [42], GZD [92], PH_VS [50], USR + GZD [84], USR + PS [84], USR + ROCS [84], GZD + PS [84], GZD + ROCS [84], PS + ROCS [84], Findsite [101], Fragsite [102], Gnina [87], GOLD-EATL [96], Glide-EATL [96], CompM [96], CompScore [75], CNN [77], DenseFS [44], SIEVE-Score [98], DeepScore [94], and RF-Score-VSv3 [98].

In particular„ we compare our methods against the well-known 3*D*-methods Ultrafast Shape Recognition (USR) [8], shape-based, ligand-centric method (ROCS) [38], PatchSurfer (PS) [42], Zernike (GZD) [92] and PH_VS [50] in Cleves-Jain dataset. In Table 1, we report the performances of all these 3*D* methods with 50 conformations [84] except PH_VS with 1 conformation [50].

In Table 2, we compare our models against other state-of-the-art VS methods on DUD-E Diverse dataset. All of these ML methods came in recent years. Among these, CNN [77] uses a convolutional neural network based framework with GPU-accelerated layer (i.e., MolGridDataLayer) to define a scoring function for protein-ligand interactions. DenseFS [44] improves the previous model CNN by a densely connected convolutional neural network architecture. Later, Sieve-Score proposes an effective scoring function: similarity of interaction energy vector score (SIEVE) [98] where they extract van der Waals, Coulomb and hydrogen bonding interactions to represent docking poses for ML. Random Forest Based Protein-ligand Scoring Function (RF-Score) [98] is another VS method proposed in the same work. Compscore [75] uses genetic algorithms to find the best combination of scoring components. Recently, energy auxiliary terms learning (EATL) [96] proposes an approach based on the scoring components extracted from the output of the scoring functions implemented in several popular docking programs like Genetic Optimisation for Ligand Docking (GOLD) [49], Molecular Operating Environment (MOE) [1] and Grid-based Ligand Docking with Energetics (GLIDE) [32, 35]. In the same work, they also combine these 3 methods and produced comprehensive EATL models, CompF and CompM. DeepScore [94] defines a deep learning based target specific scoring function. Findsite [101] proposes a threading/structure based method, where they improved it later with Fragsite [102] by using tree regression ML framework. Finally, Gnina [87] is a recently released molecular docking software, which uses deep convolutional networks to score protein-ligand structures. Note that all methods use 5-fold CV except Findsite-Fragsite (3-fold CV) and Gnina.

## C.3 Model performance across different modalities

Table 5, 6, 7, 8 show detailed ablations of the modalities used in the graph filtration step of ToDD. The success of each periodic property varies per drug target and trained ML model. However merging MP fingerprints derived from each one of these domain functions used for graph filtration has always improved the performance.

Our results also show that MP fingerprints ensure making successful predictions while training few-labeled data such as Cleves-Jain (with 2-3 labeled compounds per drug target) using a transformer-based model.

RF shows worse performance than ViT on small-scale dataset such as Cleves-Jain, despite regularization by bootstrapping and using pruned, shallow trees. Additionally, RF is more sensitive to the small variations in the training set, and imbalanced data can hamper its accuracy. In order to effectively handle the large-scale datasets that have long-tailed distributions, we undersample from the majority class (*d*ecoys). Specifically, while training RF for the binary classification task on the drug targets of DUD-E Diverse (class distributions are summarized in Table 4), we use 80% of the active compounds and the same number of randomly chosen decoys for training. Undersampling decoys to avoid heavy class imbalance achieves better trade-offs between the accuracies of active compounds and decoys.

## C.4 Computation Time

See 6.3 in the main text for details.

## C.5 Model performance using Morgan fingerprints

See 6.4 in the main text for details.

## D Societal Impact and Limitations

## D.1 Societal Impact

We perform in silico experiments and use high-throughput screening to recognize active compounds that can bind to a drug target of interest, e.g., an enzyme or protein receptor without conducting research on any human or animal subjects. Our overarching goal is to augment the capabilities of AI to enhance the in silico virtual screening and drug discovery processes, develop new drugs that have less side effects but are more effective to cure diseases, and minimize the participation of human and animal subjects as much as possible to ensure their humane and proper care.

## D.2 Limitations

We discuss in detail the computational complexity of our model in 6.3. Our model is versatile and can be scaled for large libraries by customizing the allocated computational resources. Please note that the analysis in C.4 shows the execution time of our computation pipeline when the feature extraction task is distributed across 8 cores of a single Intel Core i7 CPU. It is possible to parallelize computationally costlier operations such as VR-filtration by allocating more CPU cores on the HPC platform and optimize array operations (e.g., numpy) via the joblib library. Furthermore, all ToDD models require substantially fewer computational resources during training compared to current graph-based models that encode a compound through mining common molecular fragments, a.k.a., motifs [47]. For instance, training a motif based GNN on GuacaMol dataset which has approximately 1.5M drug-like molecules takes 130 hours of GPU time [64]. In contrast, once we generate the topological fingerprints via Vietoris-Rips filtration, training time of ToDD-ViT and ToDD-ConvNeXt for each individual drug target takes less than 1 hour on a single GPU (*N*VIDIA RTX 2080 Ti).

## Checklist

1. For all authors…
  a. Do the main claims made in the abstract and introduction accurately reflect the paper’s contributions and scope? [Yes]
  b. Did you describe the limitations of your work? [Yes] See Appendix D.
  c. Did you discuss any potential negative societal impacts of your work? [Yes] See Appendix D.
  d. Have you read the ethics review guidelines and ensured that your paper conforms to them? [Yes]
2. If you are including theoretical results…
  a. Did you state the full set of assumptions of all theoretical results? [Yes]
  b. Did you include complete proofs of all theoretical results? [Yes] Given in Appendix B.1
3. If you ran experiments…
  a. Did you include the code, data, and instructions needed to reproduce the main experimental results (either in the supplemental material or as a URL)? [Yes] Dataset links are provided. See Section 5.
  b. Did you specify all the training details (e.g., data splits, hyperparameters, how they were chosen)? [Yes] See Section 6.1
  c. Did you report error bars (e.g., with respect to the random seed after running experiments multiple times)? [No] We reported the standard deviation of our experiments evaluated by 5-fold cross-validation. See Table 1 and 2 in Section 6.2.
  d. Did you include the total amount of compute and the type of resources used (e.g., type of GPUs, internal cluster, or cloud provider)? [Yes] See Section 6.3
4. If you are using existing assets (e.g., code, data, models) or curating/releasing new assets…
  a. If your work uses existing assets, did you cite the creators? [Yes]
  b. Did you mention the license of the assets? [No] They use public-domain-equivalent license.
  c. Did you include any new assets either in the supplemental material or as a URL? [No]
  d. Did you discuss whether and how consent was obtained from people whose data you’re using/curating? [N/A]
  e. Did you discuss whether the data you are using/curating contains personally identifiable information or offensive content? [N/A]
5. If you used crowdsourcing or conducted research with human subjects…
  a. Did you include the full text of instructions given to participants and screenshots, if applicable? [N/A]
  b. Did you describe any potential participant risks, with links to Institutional Review Board (IRB) approvals, if applicable? [N/A]
  c. Did you include the estimated hourly wage paid to participants and the total amount spent on participant compensation? [N/A]

## Footnotes

* Cleves-Jain dataset: https://www.jainlab.org/Public/SF-Test-Data-DrugSpace-2006.zip

* DUD-E Diverse dataset: http://dude.docking.org/subsets/diverse

## References

[1] Molecular operating environment (moe), 2020.09 Chemical Computing Group ULC, 1010 Sherbooke St. West, Suite 910, Montreal, QC, Canada, H3A 2R7, 2022.

[2] Henry Adams and Baris Coskunuzer. Geometric approaches on persistent homology. arXiv preprint arXiv:2103.06408, 2021.

[3] Henry Adams, Tegan Emerson, Michael Kirby, Rachel Neville, Chris Peterson, Patrick Shipman, Sofya Chepushtanova, Eric Hanson, Francis Motta, and Lori Ziegelmeier. Persistence images: A stable vector representation of persistent homology. Journal of Machine Learning Research, 18, 2017.

[4] Cuneyt Gurcan Akcora, Yitao Li, Yulia R Gel, and Murat Kantarcioglu. Bitcoinheist: Topological data analysis for ransomware detection on the bitcoin blockchain. In IJCAI, 2019.

[5] Mehmet E Aktas, Esra Akbas, and Ahmed El Fatmaoui. Persistence homology of networks: methods and applications. Applied Network Science, 4(1):1–28, 2019.

[6] Erik J Amézquita, Michelle Y Quigley, Tim Ophelders, Elizabeth Munch, and Daniel H Chitwood. The shape of things to come: Topological data analysis and biology, from molecules to organisms. Developmental Dynamics, 249(7):816–833, 2020.

[7] Nieves Atienza, Rocío González-Díaz, and Manuel Soriano-Trigueros. On the stability of persistent entropy and new summary functions for topological data analysis. Pattern Recognition, 107:107509, 2020.

[8] Pedro J Ballester and W Graham Richards. Ultrafast shape recognition to search compound databases for similar molecular shapes. Journal of Computational Chemistry, 28(10):1711–1723, 2007.

[9] Andreas Bender, Hamse Y Mussa, Robert C Glen, and Stephan Reiling. Similarity searching of chemical databases using atom environment descriptors (molprint 2d): evaluation of performance. Journal of Chemical Information and Computer Sciences, 44(5):1708–1718, 2004.

[10] Nurken Berdigaliyev and Mohamad Aljofan. An overview of drug discovery and development. Future Medicinal Chemistry, 12(10):939–947, 2020.

[11] Magnus Bakke Botnan, Steffen Oppermann, and Steve Oudot. Signed barcodes for multi-parameter persistence via rank decompositions and rank-exact resolutions. arXiv preprint arXiv:2107.06800, 2021.

[12] Natasja Brooijmans and Irwin D Kuntz. Molecular recognition and docking algorithms. Annual Review of Biophysics and Biomolecular Structure, 32(1):335–373, 2003.

[13] P. Bubenik. Statistical topological data analysis using persistence landscapes. Journal of Machine Learning Research, 16(1):77–102, 2015.

[14] Zixuan Cang, Lin Mu, and Guo-Wei Wei. Representability of algebraic topology for biomolecules in machine learning based scoring and virtual screening. PLoS Computational Biology, 14(1):e1005929, 2018.

[15] Zixuan Cang and Guo-Wei Wei. Topologynet: Topology based deep convolutional and multi-task neural networks for biomolecular property predictions. PLoS Computational Biology, 13(7):e1005690, 2017.

[16] Zixuan Cang and Guo-Wei Wei. Integration of element specific persistent homology and machine learning for protein-ligand binding affinity prediction. International Journal for Numerical Methods in Biomedical Engineering, 34(2):e2914, 2018.

[17] Gunnar Carlsson. Topology and data. Bulletin of the American Mathematical Society, 46(2):255–308, 2009.

[18] Mathieu Carriere and Andrew Blumberg. Multiparameter persistence image for topological machine learning. In NeurIPS, volume 33, pages 22432–22444, 2020.

[19] Mathieu Carrière, Frédéric Chazal, Yuichi Ike, Théo Lacombe, Martin Royer, and Yuhei Umeda. Perslay: A neural network layer for persistence diagrams and new graph topological signatures. In AISTATS, pages 2786–2796, 2020.

[20] Claudio N Cavasotto and Andrew J W Orry. Ligand docking and structure-based virtual screening in drug discovery. Current Topics in Medicinal Chemistry, 7(10):1006–1014, 2007.

[21] Frédéric Chazal, Brittany Terese Fasy, Fabrizio Lecci, Alessandro Rinaldo, and Larry Wasserman. Stochastic convergence of persistence landscapes and silhouettes. In SoCG, pages 474–483, 2014.

[22] Frédéric Chazal and Bertrand Michel. An introduction to topological data analysis: fundamental and practical aspects for data scientists. Frontiers in Artificial Intelligence, 4, 2021.

[23] Yuzhou Chen, Ignacio Segovia, and Yulia R Gel. Z-GCNETs: time zigzags at graph convolutional networks for time series forecasting. In ICML, pages 1684–1694. PMLR, 2021.

[24] Yuzhou Chen, Ignacio Segovia-Dominguez, Baris Coskunuzer, and Yulia Gel. Tamp-s2gcnets: Coupling time-aware multipersistence knowledge representation with spatio-supra graph convolutional networks for time-series forecasting. In ICLR, 2022.

[25] Yu-Min Chung and Austin Lawson. Persistence curves: A canonical framework for summarizing persistence diagrams. arXiv preprint arXiv:1904.07768, 2019.

[26] Ann E Cleves and Ajay N Jain. Robust ligand-based modeling of the biological targets of known drugs. Journal of Medicinal Chemistry, 49(10):2921–2938, 2006.

[27] Tamal Krishna Dey and Yusu Wang. Computational Topology for Data Analysis. Cambridge University Press, 2022.

[28] Alexey Dosovitskiy, Lucas Beyer, Alexander Kolesnikov, Dirk Weissenborn, Xiaohua Zhai, Thomas Unterthiner, Mostafa Dehghani, Matthias Minderer, Georg Heigold, Sylvain Gelly, et al. An image is worth 16×16 words: Transformers for image recognition at scale. arXiv preprint arXiv:2010.11929, 2020.

[29] Joseph L Durant, Burton A Leland, Douglas R Henry, and James G Nourse. Reoptimization of mdl keys for use in drug discovery. Journal of Chemical Information and Computer Sciences, 42(6):1273–1280, 2002.

[30] Herbert Edelsbrunner and John Harer. Computational Topology: An Introduction. American Mathematical Society, 2010.

[31] Sean Ekins, Ana C Puhl, Kimberley M Zorn, Thomas R Lane, Daniel P Russo, Jennifer J Klein, Anthony J Hickey, and Alex M Clark. Exploiting machine learning for end-to-end drug discovery and development. Nature Materials, 18(5):435–441, 2019.

[32] Richard A Friesner, Jay L Banks, Robert B Murphy, Thomas A Halgren, Jasna J Klicic, Daniel T Mainz, Matthew P Repasky, Eric H Knoll, Mee Shelley, Jason K Perry, et al. Glide: a new approach for rapid, accurate docking and scoring. 1. method and assessment of docking accuracy. Journal of Medicinal Chemistry, 47(7):1739–1749, 2004.

[33] Marian Gidea and Yuri Katz. Topological data analysis of financial time series: Landscapes of crashes. Physica A: Statistical Mechanics and Its Applications, 491:820–834, 2018.

[34] Barbara Giunti. TDA applications library, 2022. https://www.zotero.org/groups/2425412/tda-applications/library.

[35] Thomas A Halgren, Robert B Murphy, Richard A Friesner, Hege S Beard, Leah L Frye, W Thomas Pollard, and Jay L Banks. Glide: a new approach for rapid, accurate docking and scoring. 2. enrichment factors in database screening. Journal of Medicinal Chemistry, 47(7):1750–1759, 2004.

[36] Lowell H Hall and Lemont B Kier. Electrotopological state indices for atom types: a novel combination of electronic, topological, and valence state information. Journal of Chemical Information and Computer Sciences, 35(6):1039–1045, 1995.

[37] Masahiro Hattori, Yasushi Okuno, Susumu Goto, and Minoru Kanehisa. Development of a chemical structure comparison method for integrated analysis of chemical and genomic information in the metabolic pathways. Journal of the American Chemical Society, 125(39):11853–11865, 2003.

[38] Paul CD Hawkins, A Geoffrey Skillman, and Anthony Nicholls. Comparison of shape-matching and docking as virtual screening tools. Journal of Medicinal Chemistry, 50(1):74–82, 2007.

[39] Felix Hensel, Michael Moor, and Bastian Rieck. A survey of topological machine learning methods. Frontiers in Artificial Intelligence, 4:52, 2021.

[40] Jérôme Hert, Peter Willett, David J Wilton, Pierre Acklin, Kamal Azzaoui, Edgar Jacoby, and Ansgar Schuffenhauer. New methods for ligand-based virtual screening: use of data fusion and machine learning to enhance the effectiveness of similarity searching. Journal of Chemical Information and Modeling, 46(2):462–470, 2006.

[41] Christoph Hofer, Florian Graf, Bastian Rieck, Marc Niethammer, and Roland Kwitt. Graph filtration learning. In ICML, pages 4314–4323, 2020.

[42] Bingjie Hu, Xiaolei Zhu, Lyman Monroe, Mark G Bures, and Daisuke Kihara. Pl-patchsurfer: a novel molecular local surface-based method for exploring protein-ligand interactions. International Journal of Molecular Sciences, 15(9):15122–15145, 2014.

[43] Takashi Ichinomiya, Ippei Obayashi, and Yasuaki Hiraoka. Persistent homology analysis of craze formation. Physical Review E, 95(1):012504, 2017.

[44] Fergus Imrie, Anthony R Bradley, Mihaela van der Schaar, and Charlotte M Deane. Protein family-specific models using deep neural networks and transfer learning improve virtual screening and highlight the need for more data. Journal of Chemical Information and Modeling, 58(11):2319–2330, 2018.

[45] Peiran Jiang, Ying Chi, Xiao-Shuang Li, Xiang Liu, Xian-Sheng Hua, and Kelin Xia. Molecular persistent spectral image (mol-psi) representation for machine learning models in drug design. Briefings in Bioinformatics, 23(1):bbab527, 2022.

[46] Tian Jiang, Meichen Huang, Ignacio Segovia-Dominguez, Nathaniel Newlands, and Yulia R Gel. Learning space-time crop yield patterns with zigzag persistence-based lstm: Toward more reliable digital agriculture insurance. In AAAI, volume 36, pages 12538–12544, 2022.

[47] Wengong Jin, Regina Barzilay, and Tommi Jaakkola. Hierarchical generation of molecular graphs using structural motifs. In ICML, pages 4839–4848. PMLR, 2020.

[48] Megan Johnson and Jae-Hun Jung. Instability of the betti sequence for persistent homology and a stabilized version of the betti sequence. arXiv preprint arXiv:2109.09218, 2021.

[49] Gareth Jones, Peter Willett, Robert C Glen, Andrew R Leach, and Robin Taylor. Development and validation of a genetic algorithm for flexible docking. Journal of Molecular Biology, 267(3):727–748, 1997.

[50] Bryn Keller, Michael Lesnick, and Theodore L Willke. Persistent homology for virtual screening. 2018.

[51] Michael Kerber and Alexander Rolle. Fast minimal presentations of bi-graded persistence modules. In ALENEX, pages 207–220. SIAM, 2021.

[52] Talia B Kimber, Yonghui Chen, and Andrea Volkamer. Deep learning in virtual screening: recent applications and developments. International Journal of Molecular Sciences, 22(9):4435, 2021.

[53] Justin Klekota and Frederick P Roth. Chemical substructures that enrich for biological activity. Bioinformatics, 24(21):2518–2525, 2008.

[54] Nils M Kriege, Fredrik D Johansson, and Christopher Morris. A survey on graph kernels. Applied Network Science, 5(1):1–42, 2020.

[55] Gregory Leibon, Scott Pauls, Daniel Rockmore, and Robert Savell. Topological structures in the equities market network. Proceedings of the National Academy of Sciences, 105(52):20589–20594, 2008.

[56] Christian Lemmen and Thomas Lengauer. Computational methods for the structural alignment of molecules. Journal of Computer-Aided Molecular Design, 14(3):215–232, 2000.

[57] M Lesnick. Multiparameter persistence lecture notes, 2019. https://www.albany.edu/~ML644186/AMAT_840_Spring_2019/Math840_Notes.pdf.

[58] Michael Lesnick and Matthew Wright. Computing minimal presentations and bigraded betti numbers of 2-parameter persistent homology. arXiv preprint arXiv:1902.05708, 2019.

[59] Sunhyuk Lim, Facundo Memoli, and Osman Berat Okutan. Vietoris-rips persistent homology, injective metric spaces, and the filling radius. arXiv preprint arXiv:2001.07588, 2020.

[60] Liyuan Liu, Xiaodong Liu, Jianfeng Gao, Weizhu Chen, and Jiawei Han. Understanding the difficulty of training transformers. arXiv preprint arXiv:2004.08249, 2020.

[61] Xiang Liu, Huitao Feng, Jie Wu, and Kelin Xia. Dowker complex based machine learning (dcml) models for protein-ligand binding affinity prediction. PLoS Computational Biology, 18(4):e1009943, 2022.

[62] Xiang Liu and Kelin Xia. Neighborhood complex based machine learning (ncml) models for drug design. In Interpretability of Machine Intelligence in Medical Image Computing, and Topological Data Analysis and Its Applications for Medical Data, pages 87–97. Springer, 2021.

[63] Zhuang Liu, Hanzi Mao, Chao-Yuan Wu, Christoph Feichtenhofer, Trevor Darrell, and Saining Xie. A convnet for the 2020s. arXiv preprint arXiv:2201.03545, 2022.

[64] Krzysztof Maziarz, Henry Jackson-Flux, Pashmina Cameron, Finton Sirockin, Nadine Schneider, Nikolaus Stiefl, Marwin Segler, and Marc Brockschmidt. Learning to extend molecular scaffolds with structural motifs. arXiv preprint arXiv:2103.03864, 2021.

[65] James L Melville, Edmund K Burke, and Jonathan D Hirst. Machine learning in virtual screening. Combinatorial Chemistry & High Throughput Screening, 12(4):332–343, 2009.

[66] Zhenyu Meng and Kelin Xia. Persistent spectral–based machine learning (perspect ml) for protein-ligand binding affinity prediction. Science Advances, 7(19):eabc5329, 2021.

[67] Michael M Mysinger, Michael Carchia, John J Irwin, and Brian K Shoichet. Directory of useful decoys, enhanced (dud-e): better ligands and decoys for better benchmarking. Journal of Medicinal Chemistry, 55(14):6582–6594, 2012.

[68] Takenobu Nakamura, Yasuaki Hiraoka, Akihiko Hirata, Emerson G Escolar, and Yasumasa Nishiura. Persistent homology and many-body atomic structure for medium-range order in the glass. Nanotechnology, 26(30):304001, 2015.

[69] Bruno J Neves, Rodolpho C Braga, Cleber C Melo-Filho, José Teófilo Moreira-Filho, Eugene N Muratov, and Carolina Horta Andrade. Qsar-based virtual screening: advances and applications in drug discovery. Frontiers in Pharmacology, 9:1275, 2018.

[70] Duc Duy Nguyen, Zixuan Cang, and Guo-Wei Wei. A review of mathematical representations of biomolecular data. Physical Chemistry Chemical Physics, 22(8):4343–4367, 2020.

[71] Duc Duy Nguyen, Zixuan Cang, Kedi Wu, Menglun Wang, Yin Cao, and Guo-Wei Wei. Mathematical deep learning for pose and binding affinity prediction and ranking in d3r grand challenges. Journal of Computer-Aided Molecular Design, 33(1):71–82, 2019.

[72] Duc Duy Nguyen, Kaifu Gao, Menglun Wang, and Guo-Wei Wei. Mathdl: mathematical deep learning for d3r grand challenge 4. Journal of Computer-Aided Molecular Design, 34(2):131–147, 2020.

[73] Dorcas Ofori-Boateng, I Segovia Dominguez, C Akcora, Murat Kantarcioglu, and Yulia R Gel. Topological anomaly detection in dynamic multilayer blockchain networks. In ECML PKDD, pages 788–804, 2021.

[74] Nina Otter, Mason A Porter, Ulrike Tillmann, Peter Grindrod, and Heather A Harrington. A roadmap for the computation of persistent homology. EPJ Data Science, 6:1–38, 2017.

[75] Yunierkis Perez-Castillo, Stellamaris Sotomayor-Burneo, Karina Jimenes-Vargas, Mario Gonzalez-Rodriguez, Maykel Cruz-Monteagudo, Vinicio Armijos-Jaramillo, M Natália DS Cordeiro, Fernanda Borges, Aminael Sánchez-Rodríguez, and Eduardo Tejera. Compscore: boosting structure-based virtual screening performance by incorporating docking scoring function components into consensus scoring. Journal of Chemical Information and Modeling, 59(9):3655–3666, 2019.

[76] Ralph H Petrucci, F Geoffrey Herring, and Jeffry D Madura. General Chemistry: Principles and Modern Applications. Pearson Prentice Hall, 2010.

[77] Matthew Ragoza, Joshua Hochuli, Elisa Idrobo, Jocelyn Sunseri, and David Ryan Koes. Protein–ligand scoring with convolutional neural networks. Journal of Chemical Information and Modeling, 57(4):942–957, 2017.

[78] John W Raymond, Eleanor J Gardiner, and Peter Willett. Rascal: Calculation of graph similarity using maximum common edge subgraphs. The Computer Journal, 45(6):631–644, 2002.

[79] Peter Ripphausen, Britta Nisius, and Jürgen Bajorath. State-of-the-art in ligand-based virtual screening. Drug Discovery Today, 16(9-10):372–376, 2011.

[80] David Rogers and Mathew Hahn. Extended-connectivity fingerprints. Journal of Chemical Information and Modeling, 50(5):742–754, 2010.

[81] Julian Schwartz, Mahendra Awale, and Jean-Louis Reymond. Smifp (smiles fingerprint) chemical space for virtual screening and visualization of large databases of organic molecules. Journal of Chemical Information and Modeling, 53(8):1979–1989, 2013.

[82] Chao Shen, Junjie Ding, Zhe Wang, Dongsheng Cao, Xiaoqin Ding, and Tingjun Hou. From machine learning to deep learning: Advances in scoring functions for protein–ligand docking. Wiley Interdisciplinary Reviews: Computational Molecular Science, 10(1):e1429, 2020.

[83] Zhuoran Shen, Mingyuan Zhang, Haiyu Zhao, Shuai Yi, and Hongsheng Li. Efficient attention: Attention with linear complexities. In Proceedings of the IEEE/CVF Winter Conference on Applications of Computer Vision, pages 3531–3539, 2021.

[84] Woong-Hee Shin, Xiaolei Zhu, Mark Gregory Bures, and Daisuke Kihara. Three-dimensional compound comparison methods and their application in drug discovery. Molecules, 20(7):12841–12862, 2015.

[85] Christoph Steinbeck, Christian Hoppe, Stefan Kuhn, Matteo Floris, Rajarshi Guha, and Egon L Willighagen. Recent developments of the chemistry development kit (cdk)-an open-source java library for chemo-and bioinformatics. Current Pharmaceutical Design, 12(17):2111–2120, 2006.

[86] Vladimir B Sulimov, Danil C Kutov, and Alexey V Sulimov. Advances in docking. Current Medicinal Chemistry, 26(42):7555–7580, 2019.

[87] Jocelyn Sunseri and David Ryan Koes. Virtual screening with gnina 1.0. Molecules, 26(23):7369, 2021.

[88] Dominique Sydow, Lindsey Burggraaff, Angelika Szengel, Herman WT van Vlijmen, Adriaan P IJzerman, Gerard JP van Westen, and Andrea Volkamer. Advances and challenges in computational target prediction. Journal of Chemical Information and Modeling, 59(5):1728–1742, 2019.

[89] Ashleigh Linnea Thomas. Invariants and Metrics for Multiparameter Persistent Homology. PhD thesis, Duke University, 2019.

[90] Jean-François Truchon and Christopher I Bayly. Evaluating virtual screening methods: good and bad metrics for the “early recognition” problem. Journal of Chemical Information and Modeling, 47(2):488–508, 2007.

[91] Jessica Vamathevan, Dominic Clark, Paul Czodrowski, Ian Dunham, Edgardo Ferran, George Lee, Bin Li, Anant Madabhushi, Parantu Shah, Michaela Spitzer, et al. Applications of machine learning in drug discovery and development. Nature Reviews Drug Discovery, 18(6):463–477, 2019.

[92] Vishwesh Venkatraman, Padmasini Ramji Chakravarthy, and Daisuke Kihara. Application of 3d zernike descriptors to shape-based ligand similarity searching. Journal of Cheminformatics, 1(1):1–19, 2009.

[93] Oliver Vipond. Multiparameter persistence landscapes. Journal of Machine Learning Research, 21:61–1, 2020.

[94] Dingyan Wang, Chen Cui, Xiaoyu Ding, Zhaoping Xiong, Mingyue Zheng, Xiaomin Luo, Hualiang Jiang, and Kaixian Chen. Improving the virtual screening ability of target-specific scoring functions using deep learning methods. Frontiers in Pharmacology, page 924, 2019.

[95] LIU Xiang and Kelin Xia. Persistent tor-algebra based stacking ensemble learning (pta-sel) for proteinprotein binding affinity prediction. In ICLR 2022 Workshop on Geometrical and Topological Representation Learning, 2022.

[96] Guo-Li Xiong, Wen-Ling Ye, Chao Shen, Ai-Ping Lu, Ting-Jun Hou, and Dong-Sheng Cao. Improving structure-based virtual screening performance via learning from scoring function components. Briefings in Bioinformatics, 22(3):bbaa094, 2021.

[97] Chun Wei Yap. Padel-descriptor: An open source software to calculate molecular descriptors and fingerprints. Journal of Computational Chemistry, 32(7):1466–1474, 2011.

[98] Nobuaki Yasuo and Masakazu Sekijima. Improved method of structure-based virtual screening via interaction-energy-based learning. Journal of Chemical Information and Modeling, 59(3):1050–1061, 2019.

[99] Monisha Yuvaraj, Asim K Dey, Vyacheslav Lyubchich, Yulia R Gel, and H Vincent Poor. Topological clustering of multilayer networks. Proceedings of the National Academy of Sciences, 118(21):e2019994118, 2021.

[100] Qi Zhao and Yusu Wang. Learning metrics for persistence-based summaries and applications for graph classification. In NeurIPS, volume 32, 2019.

[101] Hongyi Zhou, Hongnan Cao, and Jeffrey Skolnick. Findsitecomb2. 0: A new approach for virtual ligand screening of proteins and virtual target screening of biomolecules. Journal of Chemical Information and Modeling, 58(11):2343–2354, 2018.

[102] Hongyi Zhou, Hongnan Cao, and Jeffrey Skolnick. Fragsite: a fragment-based approach for virtual ligand screening. Journal of Chemical Information and Modeling, 61(4):2074–2089, 2021.

[103] Vincent Zoete, Antoine Daina, Christophe Bovigny, and Olivier Michielin. Swisssimilarity: a web tool for low to ultra high throughput ligand-based virtual screening. Journal of Chemical Information and Modeling, 56(8):1399–1404, 2016.

